# Transcriptome and proteome profiling reveals complex adaptations of *Candida parapsilosis* cells assimilating hydroxyaromatic carbon sources

**DOI:** 10.1101/2021.09.13.460007

**Authors:** Andrea Cillingová, Renáta Tóth, Anna Mojáková, Igor Zeman, Romana Vrzoňová, Barbara Siváková, Peter Baráth, Martina Neboháčová, Zuzana Klepcová, Filip Brázdovič, Hana Lichancová, Viktória Hodorová, Broňa Brejová, Tomáš Vinař, Ľubomír Tomáška, Atilla Gácser, Toni Gabaldón, Jozef Nosek

**Affiliations:** Department of Biochemistry, Faculty of Natural Sciences, Comenius University in Bratislava, Ilkovičova 6, 842 15 Bratislava, Slovakia; HCEMM-USZ Department of Microbiology, University of Szeged, Közép fasor 52, H-6726 Szeged, Hungary; MTA-SZTE Lendület Mycobiome Research Group, University of Szeged, Szeged, Hungary; Institute of Chemistry, Slovak Academy of Sciences, Dúbravská cesta 9, 845 38 Bratislava, Slovakia; Department of Computer Science, Faculty of Mathematics, Physics and Informatics, Comenius University in Bratislava, Bratislava, Slovakia; Department of Applied Informatics, Faculty of Mathematics, Physics and Informatics, Comenius University in Bratislava, Bratislava, Slovakia; Department of Genetics, Faculty of Natural Sciences, Comenius University in Bratislava, Ilkovičova 6, 842 15 Bratislava, Slovakia; Institute for Research in Biomedicine (IRB), The Barcelona Institute of Science and Technology, Baldiri Reixac, 10, 08028 Barcelona, Spain; Barcelona Supercomputing Centre (BSC-CNS). Jordi Girona, 29. 08034. Barcelona, Spain; Catalan Institution for Research and Advanced Studies (ICREA), Barcelona, Spain

**Author notes:** Corresponding author: (JN).

## Abstract

Many fungal species utilize hydroxyderivatives of benzene and benzoic acid as carbon sources. The yeast *Candida parapsilosis* metabolizes these compounds via the 3-oxoadipate and gentisate pathways, whose components are encoded by two metabolic gene clusters. In this study, we determine the chromosome level assembly of the *C. parapsilosis* strain CLIB214 and use it for transcriptomic and proteomic investigation of cells cultivated on hydroxyaromatic substrates. We demonstrate that the genes coding for enzymes and plasma membrane transporters involved in the 3-oxoadipate and gentisate pathways are highly upregulated and their expression is controlled in a substrate-specific manner. However, regulatory proteins involved in this process are not known. Using the knockout mutants, we show that putative transcriptional factors encoded by the genes *OTF1* and *GTF1* located within these gene clusters function as transcriptional activators of the 3-oxoadipate and gentisate pathway, respectively. We also show that the activation of both pathways is accompanied by upregulation of genes for the enzymes involved in β-oxidation of fatty acids, glyoxylate cycle, amino acid metabolism, and peroxisome biogenesis. Transcriptome and proteome profiles of the cells grown on 4-hydroxybenzoate and 3-hydroxybenzoate, which are metabolized via the 3-oxoadipate and gentisate pathway, respectively, reflect their different connection to central metabolism. Yet we find that the expression profiles differ also in the cells assimilating 4-hydroxybenzoate and hydroquinone, which are both metabolized in the same pathway. This finding is consistent with the phenotype of the Otf1p-lacking mutant, which exhibits impaired growth on hydroxybenzoates, but still utilizes hydroxybenzenes, thus indicating that additional, yet unidentified transcription factor could be involved in the 3-oxoadipate pathway regulation. Moreover, we propose that bicarbonate ions resulting from decarboxylation of hydroxybenzoates also contribute to differences in the cell responses to hydroxybenzoates and hydroxybenzenes. Finally, our phylogenetic analysis highlights evolutionary paths leading to metabolic adaptations of yeast cells assimilating hydroxyaromatic substrates.

**Author summary:** Benzene and its derivatives are simple aromatic compounds representing key substances for the chemical industry. While benzene itself is toxic and carcinogenic, benzoic acid is commonly used in the food industry and some of its derivatives are used in pharmacology (aspirin) or cosmetics (parabens). The benzene ring of aromatic molecules is relatively stable, but many microorganisms including yeasts break it enzymatically and, in a series of biochemical reactions, utilize resulting metabolites as carbon sources. Understanding the genetic basis of corresponding metabolic pathways and their regulation opens a venue for applications in biotechnology and bioremediation of polluted environments. Here we investigate the yeast *Candida parapsilosis* which assimilates various hydroxybenzenes and hydroxybenzoates via the 3-oxoadipate and gentisate pathways. We show that the genes coding for the substrate transporters and enzymes involved in both pathways are co-expressed and regulated by the transcriptional activators Otf1p and Gtf1p, respectively. Our results also reveal the connections of both pathways to central metabolism and organelle biogenesis and provide an insight into evolution of metabolism of hydroxyaromatic compounds.

## Introduction

Metabolic gene clusters (MGCs) are composed of co-localized genes, whose products participate in the same metabolic pathway. In most cases, their functions are linked to the production of secondary metabolites or the assimilation of unconventional substrates. Such biochemical pathways are usually nonessential, but in specific circumstances they may provide a growth benefit for the host organism. In general, MGCs encode the enzymes catalyzing reactions in a biochemical pathway, membrane transporters for substrates or metabolites, as well as transcription factors that control the expression of corresponding genes. Gene clustering thus generates functional genetic modules whose co-regulated expression facilitates rapid adaptation of cellular metabolism to environmental changes (1, 2). The occurrence of MGCs in eukaryotic genomes was originally considered to be rare. However, bioinformatic analyses of a constantly increasing number of sequenced genomes show that the gene clusters are their typical feature, especially in case of fungal and plant genomes (3–5). The formation of MGCs also facilitates their transmission via horizontal gene transfer, thus contributing to metabolic diversity of fungal species and their ecological adaptation (6). Investigations of MGCs provide a venue for elucidating their evolutionary origin, genetic organization, and expression, as well as the coordination of the corresponding biochemical pathways with the central cellular metabolism.

Previously, we identified and characterized several genes from the pathogenic yeast *Candida parapsilosis* arranged in two MGCs, which are conserved in the genomes of yeast species from the ‘CUG-Ser1’ clade of the subphylum Saccharomycotina (7–12). These MGCs code for enzymes of the gentisate (GP) and 3-oxoadipate (3-OAP) pathways that are involved in catabolic degradation of a broad spectrum of hydroxyderivatives of benzene and benzoic acid. While 3-hydroxybenzoate and gentisate (2,5-dihydroxybenzoate) are metabolized via the GP, 4-hydroxybenzoate, 2,4-dihydroxybenzoate, protocatechuate (3,4-dihydroxybenzoate), hydroquinone, and resorcinol are degraded via the hydroxyhydroquinone (HHQ) branch of the 3-OAP (13, 14). The resulting products of both biochemical pathways (i.e. fumarate and pyruvate in the GP; succinate and acetyl-CoA in the 3-OAP) can be channeled into tricarboxylic acid (TCA) cycle operating in mitochondria. Interconnection of both pathways with these organelles is mediated by metabolite carriers in the inner mitochondrial membrane (i.e. Sfc1p, Leu5p, Yhm2p, and Mpc1p). Moreover, the enzymes catalyzing the last two steps of the 3-OAP (i.e. 3-oxoadipate:succinyl-CoA transferase (Osc1p) and 3-oxoadipyl-CoA thiolase (Oct1p)) are imported into mitochondria (9, 11). In addition, we have previously identified a family of genes coding for the plasma membrane transporters for hydroxybenzoates (10). Moreover, while both pathways are repressed in cells assimilating glucose, corresponding genes are highly induced during cultivation on media containing a hydroxyaromatic substrate as a sole carbon source (7,9–11). Although the transcriptional factors involved in this regulation have not yet been identified, both MGCs contain a gene for uncharacterized zinc cluster transcription factor representing a candidate transcriptional activator of the corresponding pathway.

In this study, we investigate the regulation of the 3-OAP and GP as well as the coordination of both pathways with central metabolism and organelle biogenesis. Using the analysis of transcriptomic and proteomic profiles of *C. parapsilosis* cells assimilating hydroxyaromatic compounds we show that the induction of both pathways is accompanied by the upregulation of genes whose products are involved in β-oxidation of fatty acids (FA), glyoxylate cycle, metabolism of amino acids, and the biogenesis of peroxisomes. Our results also highlight the differences between the metabolism of hydroxybenzoates and hydroxybenzenes. Moreover, we demonstrate experimentally that putative transcription factors named Gtf1p and Otf1p function as transcriptional activators of the GP and 3-OAP genes, respectively. Their phylogenetic analysis shed additional insight into the evolution of both biochemical pathways.

## Results and Discussion

### Gene expression landscape of *C. parapsilosis* cells assimilating hydroxyaromatic carbon sources

Several studies have demonstrated that *C. parapsilosis* assimilates a broad spectrum of hydroxyderivatives of benzene and benzoic acid via the GP and 3-OAP (7, 13). To investigate the regulation of both pathways we analyzed *C. parapsilosis* cells grown in media containing hydroxyaromatic compounds degraded either via the 3-OAP or GP. The gene expression analysis was performed in the strain CLIB214 (CBS604), which is together with derived mutants commonly used in experimental studies (e.g. 15-18). This strain was originally isolated from a patient with tropical diarrhea in Puerto Rico (19) and it represents the type strain of *C. parapsilosis*. Although a genome sequence survey of CLIB214 was carried out in 2005 by Sanger sequencing (20), the complete genome sequence of this strain was not available. Here, we determined the chromosome level genome assembly of the CLIB214 strain by combining Oxford Nanopore and Illumina sequencing technologies and used it for analyses of cells utilizing hydroxyaromatic substrates (see below). The resulting CLIB214 assembly has a total length of 13.0 Mbp and consists of 8 nuclear chromosomes corresponding to the electrophoretic karyotype determined by PFGE (**Fig 1**). Alignments with the reference genome sequence of the strain CDC317 (21) cover 99.5% of the assembly and have a 99.9% identity. Compared to the CDC317 assembly, there is a single large-scale translocation between chromosomes 4 and 5 (CDC317 contigs HE605208.1 and HE605204.1). Annotation of the nuclear chromosomes contains 5,856 predicted protein-coding genes; 5,797 of them overlap with protein coding genes mapped from the CDC317 strain. The six genes coding for the GP components (i.e. *MNX2, HBT1*, *GDX1*, *FPH1*, *GFA1, GTF1*) are localized in a single MGC which is present in the subtelomeric region of chromosome 6. The 3-OAP components are encoded by a cluster comprising four genes (i.e. *FRD1*, *HDX1*, *OSC1, OTF1*) located on chromosome 5, as well as by several additional loci on chromosomes 1 and 2 (**Fig 1)**.

**Fig 1.**
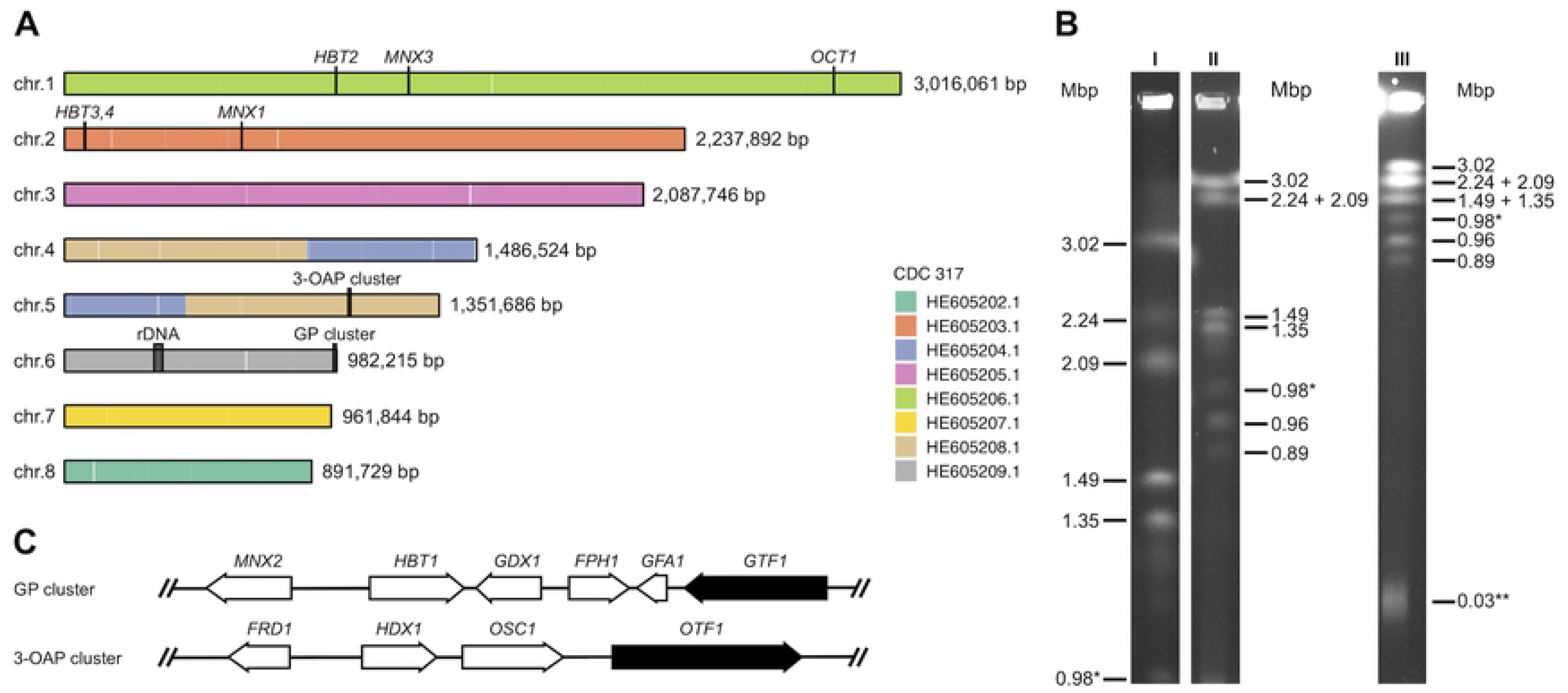
Nuclear genome organization of the *C. parapsilosis* strain CLIB214. (A) Chromosomal contigs of *C. parapsilosis* CLIB214. The colouring is based on alignments with the nuclear contigs of the reference genome sequence (CDC317) (see Materials and Methods for details). (B) Electrophoretic karyotype of CLIB214. DNA samples prepared in agarose blocks were separated by PFGE at three different conditions (I, II, and III) as described in Materials and Methods. The bands corresponding to the chromosome containing an rDNA array (0.98 Mbp) and the linear mitochondrial DNA (32.8 kbp) are indicated by one and two asterisks, respectively. (C) Organization of the GP and 3-OAP gene clusters. The genes coding for the transcription activators Gtf1p and Otf1p investigated in this study are shown in black.

Next, we used the CLIB214 genome assembly as a reference for transcriptomic and proteomic experiments to investigate the activation of genes involved in the 3-OAP and GP and their links to central cellular metabolism and organelle biogenesis. In these experiments, we compared CLIB214 cells assimilating 4-hydroxybenzoate, hydroquinone (both metabolized via the 3-OAP) and 3-hydroxybenzoate (metabolized via the GP), with those utilizing galactose as a control carbon source. By RNA-Seq analysis, we identified 270, 435, and 365 genes upregulated more than four-fold in cells cultivated in media containing 3-hydroxybenzoate, 4-hydroxybenzoate, and hydroquinone, respectively, compared to control cells grown on galactose (**Fig 2**, **S1 Table)**.

**Fig 2.**
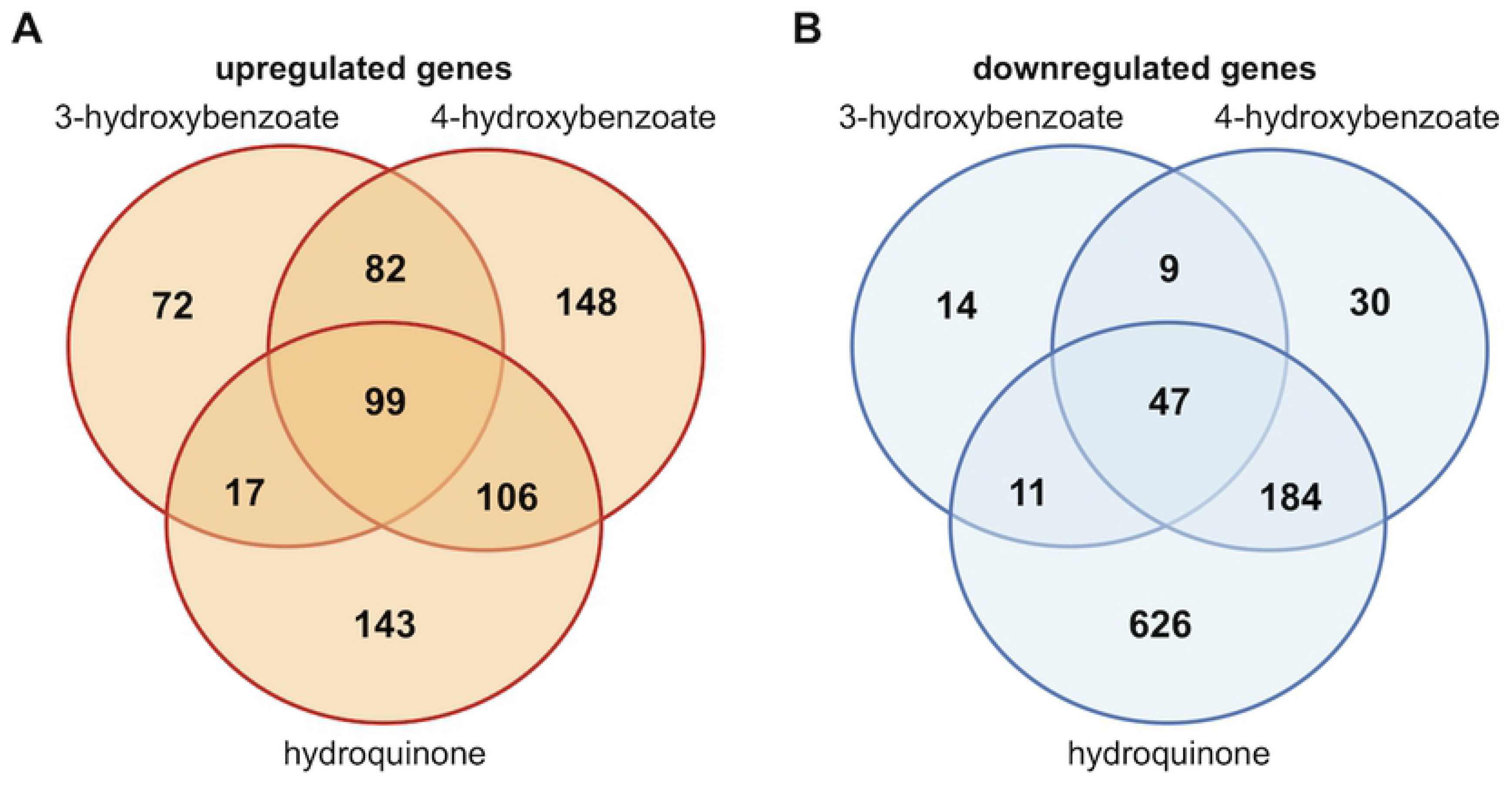
Differentially expressed genes identified by RNA-Seq analysis. The Venn diagrams show numbers of upregulated (log_2_ fold change ≥ 2; p ≤ 0.05; (A)) or downregulated (log_2_ fold change ≤ −2; p ≤ 0.05; (B)) genes in CLIB214 cells assimilating 3-hydroxybenzoate, 4-hydroxybenzoate or hydroquinone compared to galactose. The results are based on the lists of differentially expressed genes (**S1 Table**). The diagrams were drawn with a web tool (http://bioinformatics.psb.ugent.be/webtools/Venn/).

In line with our previous reports (7,9,10), the RNA-Seq analysis showed that the genes encoding the enzymes catalyzing reactions in each pathway as well as the plasma membrane carriers facilitating the transport of hydroxybenzoates (**Fig 3A**) are co-regulated in a substrate-specific manner. Specifically, the GP cluster genes are highly upregulated (i.e. between 267-(*GFA1*) and 3,061-fold (*HBT1*)) in the cells assimilating 3-hydroxybenzoate, which is metabolized via the GP. These genes exhibit only minor changes in media containing 4-hydroxybenzoate, except for *GTF1* and *HBT1* showing about 12.6- and 4.7-fold induction on this substrate, respectively (**Fig 3B**, **S1 Table**). The genes for the 3-OAP enzymes and two plasma membrane transporters (*HBT2* and its paralog *HBT3*) are highly upregulated on both 4-hydroxybenzoate (i.e. between 46.5- (*OSC1*) to 1,090-fold (*HBT2*)) and hydroquinone (i.e. between 8.1- (*HBT3*) and 208-fold (*HDX1*)). Expression of these genes changes only slightly on the GP substrate, except *MNX1* which exhibits about 19-fold increase (**Fig 3B**, **S1 Table**).

**Fig 3.**
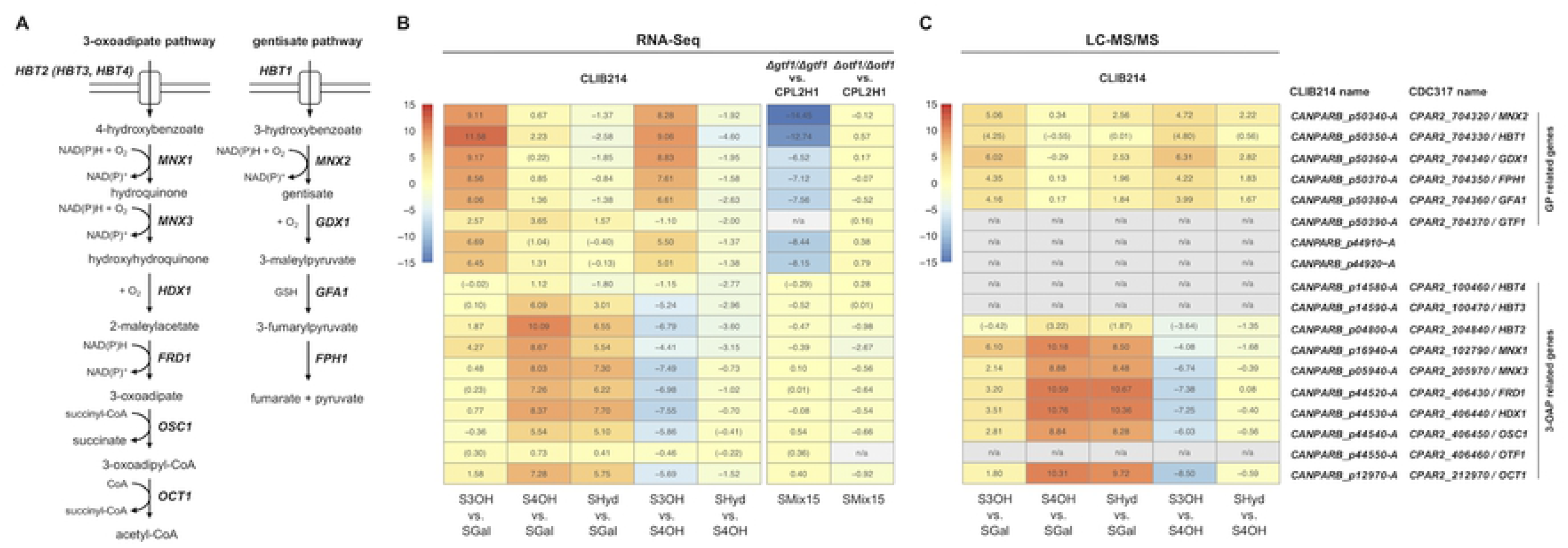
The 3-OAP and GP are induced in *C. parapsilosis* cells assimilating hydroxyaromatic compounds. (A) The simplified schemes depicting the enzymes and hydroxybenzoate transporters involved in the 3-OAP and GP in *C. parapsilosis*. (B) Differential expression of selected *C. parapsilosis* genes involved in the metabolism of hydroxyaromatic compounds. The expression was analyzed in CLIB214 cells grown to OD_600_ ∼ 1 in synthetic media containing 3-hydroxybenzoate, 4-hydroxybenzoate or hydroquinone as a sole carbon source compared to the cells cultivated in medium with galactose (i.e. S3OH vs. SGal, S4OH vs. SGal, SHyd vs. SGal) or 4-hydroxybenzoate (S3OH vs. S4OH, SHyd vs. S4OH). Analysis of the mutants Δ*gtf1/Δgtf1* and Δ*otf1/Δotf1* is based on the comparison to the parental strain CPL2H1 (Δ*gtf1/Δgtf1* vs. CPL2H1 and Δ*otf1/Δotf1* vs. CPL2H1) grown in an SMix15 medium containing three hydroxyaromatic carbon sources (i.e. 3-hydroxybenzoate, 4-hydroxybenzoate, and hydroquinone). The log_2_ fold change values are shown (**S1 Table** and **S2 Table**). Note that the values that are not statistically significant (i.e. p > 0.05) are shown in parentheses. (C) LC-MS/MS analysis of protein extracts from *C. parapsilosis* CLIB214. The cells were pre-cultivated overnight in an S3OH medium, inoculated to SGal, S3OH, S4OH, and SHyd media, and grown to ∼ 10^7^ cells/ml. Soluble proteins were extracted and analyzed by LC-MS/MS. Log2 values of mean LFQ intensity ratios are shown (**S3 Table**). For proteins that were not identified on all carbon sources the LFQ values imputed from a normal distribution were used in the calculation (indicated in parentheses).

Next, we analyzed the proteins in the cellular extracts prepared from the CLIB214 cultures by LC-MS/MS. In total, we identified 1451 proteins, of which 1176 had significantly different relative abundance (based on LFQ values) as evaluated by ANOVA test. The comparison of the 3-OAP and GP related proteins identified in the cells assimilating hydroxyaromatic substrates with those utilizing galactose shows a pattern similar to the RNA-Seq results, i.e. the 3-OAP enzymes are highly enriched on both 4-hydroxybenzoate and hydroquinone, and the GP enzymes are enriched on 3-hydroxybenzoate (**Fig 3C**, **S3 Table**). However, we did not identify several proteins (i.e. Hbt3p, Hbt4p, Gtf1p, Otf1p). We presume that this is caused by overall low abundance of these polypeptides in the cells or their depletion from the prepared extracts due to insolubility or subcellular localization.

The MGCs contain yet uncharacterized genes *FRD1* and *GFA1* which are highly induced on hydroxyaromatic substrates and appear to be co-regulated with the genes for the 3-OAP or GP enzymes, respectively. This indicates that their products could participate in the metabolism of hydroxyaromatic substrates. Based on the expression profiles and identified protein domains, we hypothesize that *FRD1* (flavin re<u>d</u>uctase 1; *CANPARB_p44520-A* (*CPAR2_406430* in CDC317)) and *GFA1* (<u>g</u>lutathione-dependent formaldehyde-<u>a</u>ctivating enzyme 1; *CANPARB_p50380-A* (*CPAR2_704360* in CDC317)) code for maleylacetate reductase and glutathione-dependent maleylpyruvate isomerase involved in the 3-OAP and GP, respectively. Moreover, the transcriptome analysis also revealed that two neighboring open reading frames (ORFs) *CANPARB_p44920-A* and *CANPARB_p44910-A* also belong to highly upregulated genes (i.e. 83- and 103-fold, respectively) in CLIB214 cells assimilating 3-hydroxybenzoate compared to galactose. These ORFs are not annotated in the reference genome (CDC317), although their sequences are identical in both strains. The deduced amino acid sequences of CANPARB_p44920-A and CANPARB_p44910-A are highly similar to the N- and C-terminal half, respectively, of bacterial proteins from the amidohydrolase superfamily, which includes 2-amino-3-carboxymuconate-6-semialdehyde decarboxylase (ACMSD), orsellinate decarboxylase (OrsB), 6-methylsalicylate decarboxylase (YanB), and salicylate decarboxylase involved in metabolism of various hydroxyaromatic compounds (**S1 Fig**, see below for a more detailed discussion).

### Metabolic pathways activated in cells assimilating hydroxyaromatic compounds

Gene ontology (GO) analysis revealed that the lists of upregulated genes in the RNA-Seq experiment are enriched for categories annotated as oxidoreductase (13.7%), hydrolase (10.7%), transferase (9.0%), transporter (8.2%), DNA- (6.1%), and protein-binding (5.8%) activities. The KEGG mapper analysis identified 163 upregulated genes coding for enzymes involved in metabolic pathways (**Fig 4**, **S2 Fig**). A large proportion of these enzymes are involved in FA metabolism (22) including those involved in β-oxidation and lipases involved in mobilization of FAs from mono- and triglycerides (MAG and TAG lipases) and (lyso)phospholipids (phospholipases B/A2). The supposedly increased level of hydrogen peroxide in peroxisomes is accompanied by overexpression of the genes for catalases, superoxide dismutase, and glutathione-*S*-transferases. Although the localization of these proteins in peroxisomes was not tested experimentally, it can be supposed that the catalase isoenzymes play a protective role in β-oxidation (23). Furthermore, the β-oxidation cycle is provided by acyl-CoA by the action of fatty acid-CoA synthetases (FAAs) that constitute an unusually large family of isoenzymes in *C. parapsilosis*. In *Saccharomyces cerevisiae,* there are four FAAs with various roles in FA metabolism, transport, acylation of proteins, vesicular transport, and transcription regulation (24). *C. albicans*, *C. auris*, *C. dubliniensis*, and *C. glabrata* contain up to five FAAs, whereas *C. parapsilosis* contains twelve FAA-encoding genes. Nine of them are highly upregulated (log_2_ fold change > 2) on at least one substrate and five on all three tested hydroxyaromatic carbon sources (**S3 Fig**). Acetyl-CoA resulting from β-oxidation is feeding downstream metabolic processes including glyoxylate cycle. Indeed, the genes for citrate synthase (*CIT1*), isocitrate lyase (*ICL1*), and malate synthase (*MLS1*) are upregulated and so CoA can be provided back to β-oxidation (25). The gene encoding a peroxisomal coenzyme A diphosphatase (*PCD1*) regenerating CoA within peroxisomes (26) is also upregulated. In addition, the genes encoding enzymes involved in carnitine shuttle, such as *CAT2* encoding a homolog of a major form of carnitine acetyltransferase with dual localization to mitochondria and peroxisomes, are upregulated supplying a shuttle of acetyl units between these organelles (27). Finally, the genes for numerous enzymes involved in metabolism of amino acids, vitamins, purines, and pyrimidines are also overexpressed thus contributing to the metabolic needs of the cells utilizing hydroxyaromatic substrates (**S2 Fig**). In many cases, the gene expression profiles based on the RNA-Seq experiment more or less correspond to those obtained by the LC-MS/MS analysis. However, there are several notable differences. In particular, the genes coding for the three glyoxylate cycle enzymes, namely citrate synthase (Cit1p), isocitrate lyase (Icl1p), and malate synthase (Mls1p) exhibit upregulated transcription (log2 fold change ≥ 2) both on 3-hydroxybenzoate and 4-hydroxybenzoate, yet the LC-MS/MS analysis indicates a slight decrease of the corresponding proteins on the former substrate. Discordances between mRNA and protein levels are usually caused by posttranscriptional regulation of protein synthesis and/or degradation (28, 29). The observed differences in transcriptome and proteome profiles imply that the interconnection of final products of the GP and 3-OAP with the intermediate metabolism differs. The last step of the GP occurs in cytosol (9) producing pyruvate and fumarate. The former could be carboxylated to oxaloacetate (30) thus supplying the substrate for sugar synthesis in the same compartment. On the other hand, the 3-OAP producing succinate and acetyl-CoA in mitochondria (11) needs the peroxisomal glyoxylate cycle to convert the latter C2 unit to C4 to supply the gluconeogenesis with a C4 substrate (31).

**Fig 4.**
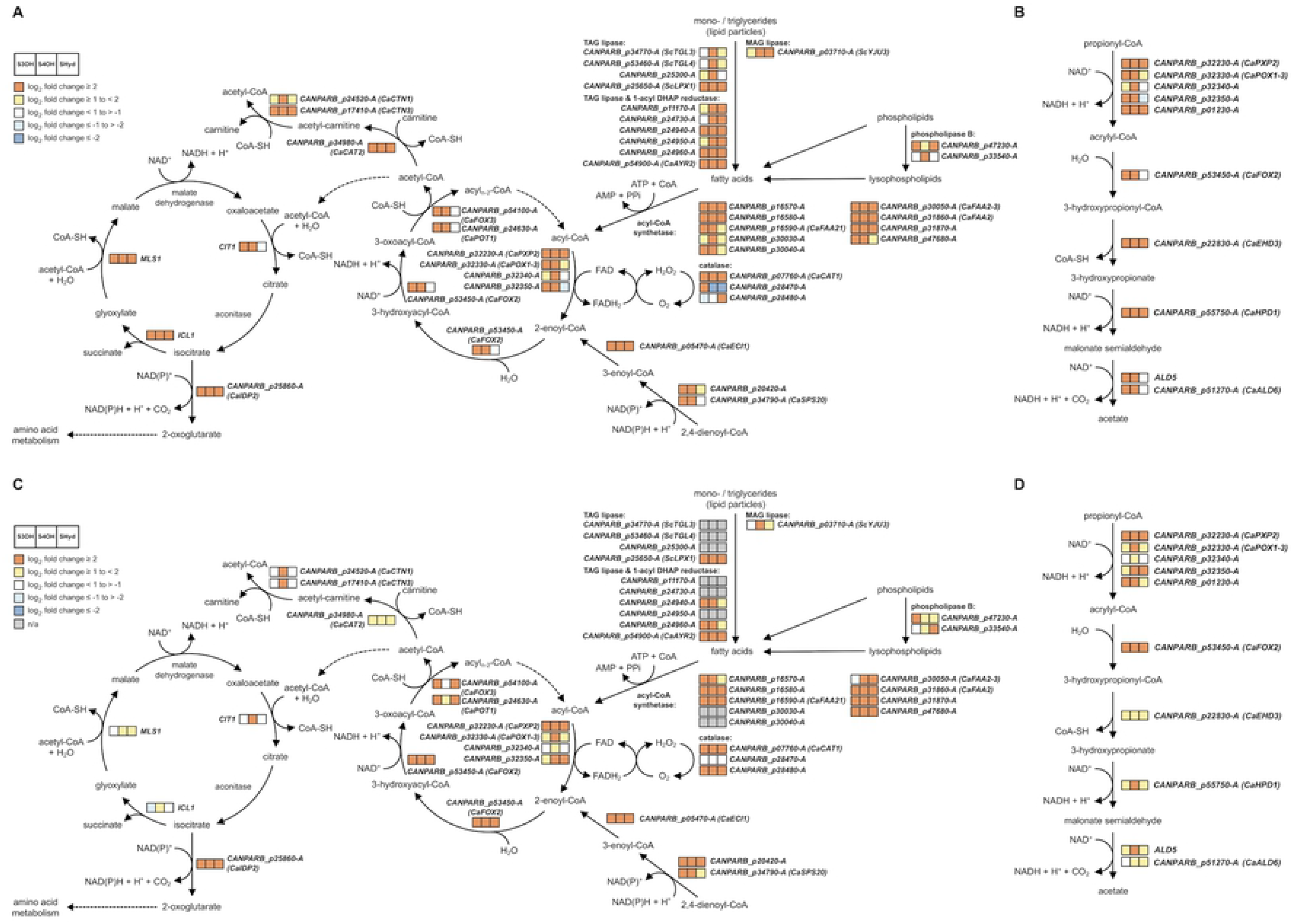
Major metabolic pathways upregulated in the cells utilizing 3-hydroxybenzoate, 4-hydroxybenzoate or hydroquinone as a sole carbon source. (A,C) The glyoxylate cycle, β-oxidation, and (B,D) modified β-oxidation pathway (32) are depicted in a simplified form. The expression profiles obtained by the RNA-Seq (A,B; **S1 Table**) and LC-MS/MS analyses (C,D; **S3 Table**) are shown. The three squares illustrate the gene expression changes on different hydroxyaromatic substrates compared to galactose as indicated in the legend in the upper left corner on panels (A) and (C). Only the genes whose transcription was overexpressed on at least one hydroxyaromatic substrate are shown. Note that not all enzymes listed in **S2 Fig** and **S4 Fig** are shown on the scheme.

### Catabolism of hydroxyaromatic compounds leads to upregulation of genes involved in the biogenesis and metabolism of peroxisomes

Previously, we reported that catabolism of hydroxyaromatic compounds is linked to mitochondria (9). Here we show that peroxisomes also play a role in cellular response to these substrates. These organelles are highly dynamic and tightly regulated by processes of *de novo* formation, division, and autophagic degradation. In yeast cells, their number depends on the utilized carbon source (33). The fact that boosting FA catabolism is accompanied by proliferation of peroxisomes in *C. parapsilosis* is underlined not only by upregulation of genes for metabolic enzymes, but also those involved in peroxisome biogenesis (**Fig 5A**; **S4 Fig**) including Pex11p crucial for peroxisome proliferation (34, 35), Pex3p and Pex19p essential for the formation of peroxisomal membrane (36), receptors Pex5p and Pex7p, and other components of matrix protein importomer, namely Pex1p, Pex2p, Pex4p, Pex8p, Pex10p, Pex12p-Pex14p, and Pex17p (37). In addition, the gene coding for inheritance protein Inp1p which secures a balanced distribution of peroxisomes between mother and daughter cells is also upregulated (38). To demonstrate the presence of peroxisomes in cells assimilating the 3-OAP and GP substrates, we constructed the plasmid pBP7-mCherry-SKL expressing a soluble codon-optimized mCherry protein (39) tagged with peroxisomal targeting signal type 1 (PTS1) serine–lysine–leucine (SKL) at its C-terminus. The plasmid pBP7-mCherry expressing an unmodified marker was used as a control. Both plasmids were introduced into *C. parapsilosis* CDU1 cells and the transformants were grown in synthetic media containing galactose. Cells containing pBP7-mCherry-SKL were also cultivated in synthetic media containing 3-hydroxybenzoate or 4-hydroxybenzoate as a sole carbon source. Examination of the transformants by fluorescence microscopy showed the presence of multiple bright foci in the cells expressing mCherry-SKL protein. The control cells carrying the pBP7-mCherry plasmid show cytosolic localization of the marker (**Fig 5B**). This result indicates that cells utilizing hydroxybenzoate substrates metabolized via the 3-OAP or GP contain multiple peroxisomes, although their number and size in the cells grown on hydroxybenzoate substrates and galactose do not seem to be substantially different.

**Fig 5.**
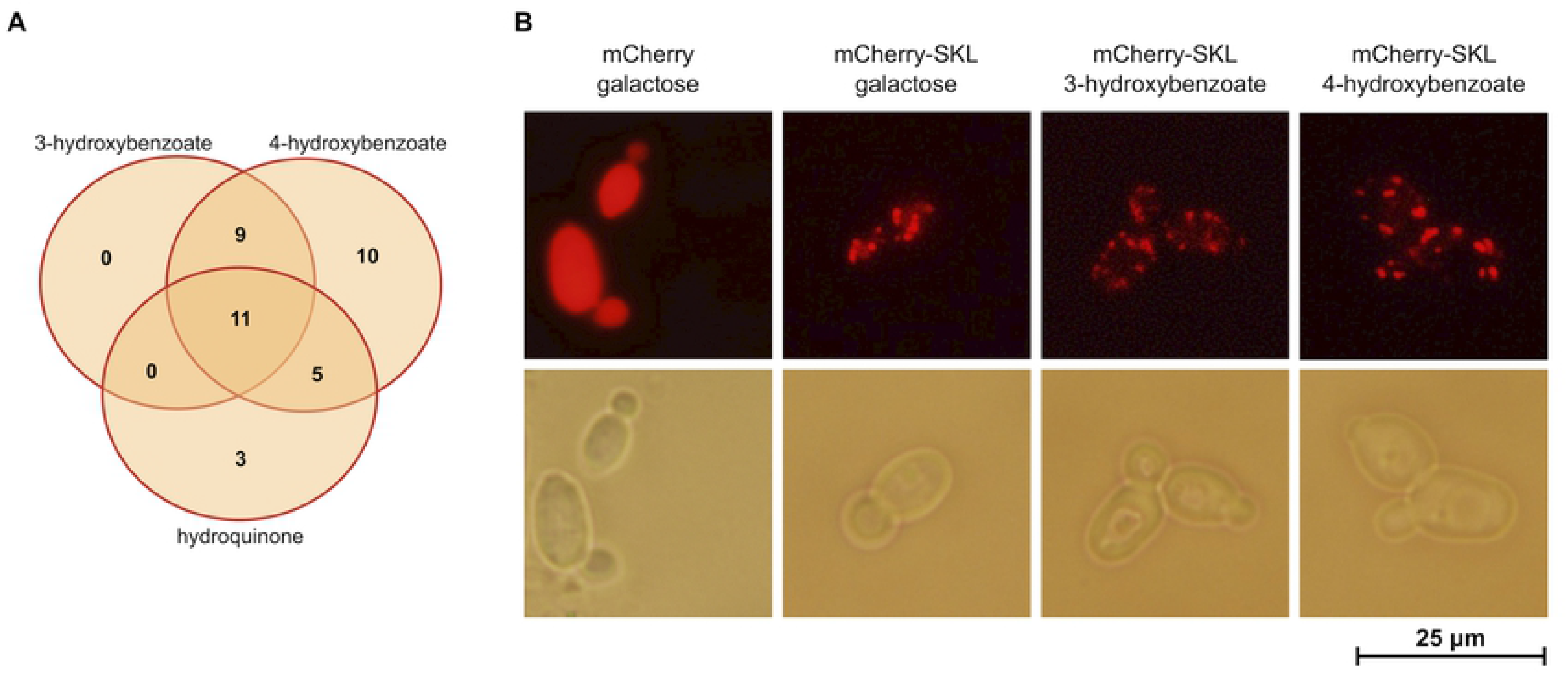
Peroxisomes are involved in the catabolism of hydroxyaromatic compounds. (A) The Venn diagram illustrating numbers of upregulated genes involved in the biogenesis and metabolism of peroxisomes identified by RNA-Seq analysis of CLIB214 cells assimilating 3-hydroxybenzoate, 4-hydroxybenzoate or hydroquinone compared to galactose. The upregulated genes (log_2_ fold change ≥ 2; p ≤ 0.05) classified into categories ‘peroxisome’, ‘peroxisomal matrix’, ‘peroxisomal membrane’ or ‘peroxisomal importomer complex’ (based on the GO subcellular component analysis; http://www.candidagenome.org) were selected from the lists of differentially expressed genes (**S1 Table; S4 Fig**). The diagram was drawn with a web tool (http://bioinformatics.psb.ugent.be/webtools/Venn/). (B) *C. parapsilosis* CDU1 cells expressing cytosolic (mCherry) and peroxisomal (mCherry-SKL) versions of the marker protein. The cells were transformed with pBP7-mCherry or pBP7-mCherry-SKL plasmids and the transformants were cultivated on a synthetic medium with galactose, 3-hydroxybenzoate or 4-hydroxybenzoate at 28 °C.

### *C. parapsilosis* response to hydroxybenzenes and hydroxybenzoates

The size and morphology of the *C. parapsilosis* colonies grown in synthetic media indicate that the cells respond differently to assimilated hydroxyaromatic substrates (**Fig 6A**). The RNA-Seq analysis of the cells utilizing 3-hydroxybenzoate, 4-hydroxybenzoate or hydroquinone show that although there is a group of ninety nine genes upregulated on any of the three carbon sources, many genes are selectively induced only on a single substrate (**Fig 2A**). As 3-hydroxybenzoate and 4-hydroxybenzoate are catabolized by distinct biochemical pathways producing different metabolites (i.e. acetyl-CoA and succinate in the 3-OAP, fumarate and pyruvate in the GP), the differences in transcription profiles of cells utilizing these substrates may reflect, at least in part, different links of these pathways to central metabolism. However, 4-hydroxybenzoate and hydroquinone are degraded in the same pathway (i.e. 3-OAP), yet only about a half of the upregulated genes are induced on both substrates and the difference in the lists of downregulated genes on these substrates is even greater (**Fig 2**). In the 3-OAP, hydroxybenzoates are decarboxylated to hydroxybenzenes (i.e. 4-hydroxybenzoate to hydroquinone; 2,4-dihydroxybenzoate and protocatechuate to hydroxyhydroquinone). The decarboxylation step is catalyzed by the monooxygenase Mnx1p which has broad substrate specificity (40, 41). This reaction releases a molecule of carbon dioxide, which can be readily converted by carbonic anhydrase to a bicarbonate anion (HCO_3_^-^). To monitor the formation of bicarbonate anions we cultivated CLIB214 cells in synthetic media containing various carbon sources and a pH indicator (bromothymol blue, p*K*a=7). We observed dramatic pH changes in the cultures grown on hydroxybenzenes compared to those assimilating hydroxybenzoates. As judged from the color of the pH indicator observed at later cultivation stages (> 12 hours), the media were acidified when the cells assimilated hydroquinone or resorcinol, which is also typical for sugar utilization (42). In contrast, the cells utilizing hydroxybenzoates (i.e. 4-hydroxybenzoate, protocatechuate, 3-hydroxybenzoate, gentisate) alkalinized the medium pointing to a buffering effect of generated bicarbonate anions (**Fig 6B**). As these ions have a role in intracellular signaling (via activation of adenylyl cyclase) and the control of metabolism (43), our results indicate that assimilation of hydroxyaromatic substrates is accompanied by a complex cellular response which is dependent on particular carbon source.

**Fig 6.**
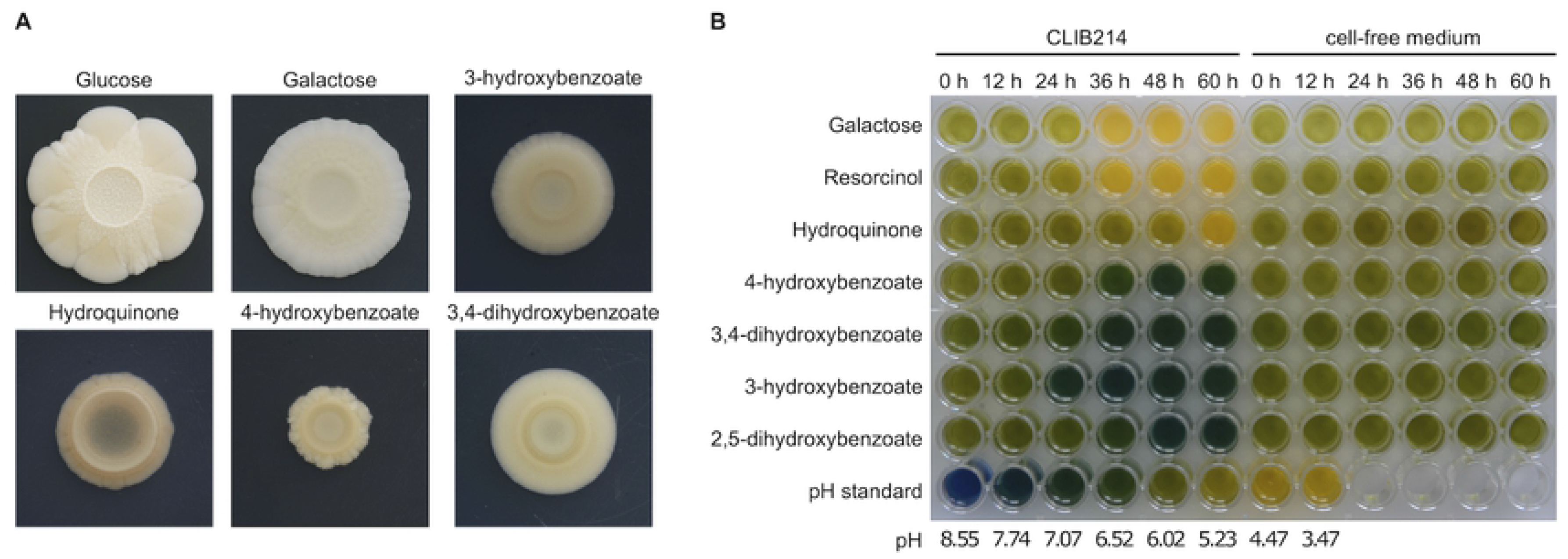
Colony morphology and pH changes in cultivation media. (A) Morphology of CLIB214 colonies grown in synthetic media differing by the carbon source. The cells were pre-grown in a complex medium (YPD) for 24 h at 28°C, washed with water, resuspended to ∼10^7^ cells/ml and 40 µl aliquots were spotted onto Petri plates containing synthetic media with indicated carbon source. The plates were incubated for 30 days at 28°C. (B) CLIB214 cells assimilating hydroxyaromatic substrates alter the pH of cultivation media. The cells were cultivated in liquid synthetic media containing indicated carbon source and bromothymol blue as pH indicator (see Materials and Methods for details). The cultures were grown for up to 60 hours at 28 °C. The cells were removed by centrifugation and the cultivation media as well as the cell-free controls were transferred into wells of a 96-well plate and photographed.

### *OTF1* and *GTF1* code for Zn(II)_2_Cys_6_ transcription activators involved in the control of the 3-OAP and GP genes, respectively

As mentioned above the 3-OAP and GP gene clusters contain the genes *OTF1* and *GTF1*, respectively, coding for putative transcription factors. The predicted proteins are 963 and 741 amino acids long, respectively, and contain Gal4-like Zn(II)_2_Cys_6_ zinc cluster DNA-binding and fungal transcription factor domains as well as putative nuclear localization signals (NLS) indicating their import into the cell nucleus (**S5 Fig**). As the orthologs of these genes are conserved in several species belonging to the ‘CUG-Ser1’ clade (see below) which assimilate hydroxyaromatic compounds (7–9) we hypothesized that *OTF1* (3-<u>o</u>xoadipate pathway transcription factor 1; *CANPARB_p44550-A* (*CPAR2_406460* in CDC317)) and *GTF1* (<u>g</u>entisate pathway transcription factor 1; *CANPARB_p50390-A* (*CPAR2_704370* in CDC317)) control the expression of corresponding MGC.

First, we confirmed that Otf1p and Gtf1p are targeted into the cell nucleus. We prepared the plasmid constructs expressing these proteins tagged with yEGFP3 at their N-termini (i.e. yEGFP3-Otf1p, yEGFP3-Gtf1p) in *C. parapsilosis* SR23 met1^-^ cells. Examination by fluorescence microscopy showed that both proteins co-localize with DAPI-stained nuclear DNA. Moreover, yEGFP3-Gtf1p appears to be concentrated in distinct foci pointing to its specific subnuclear localization. We presume that it associates with the promoters of the GP cluster genes present in the subtelomeric region of chromosome 6 (**Fig 7**).

**Fig 7.**
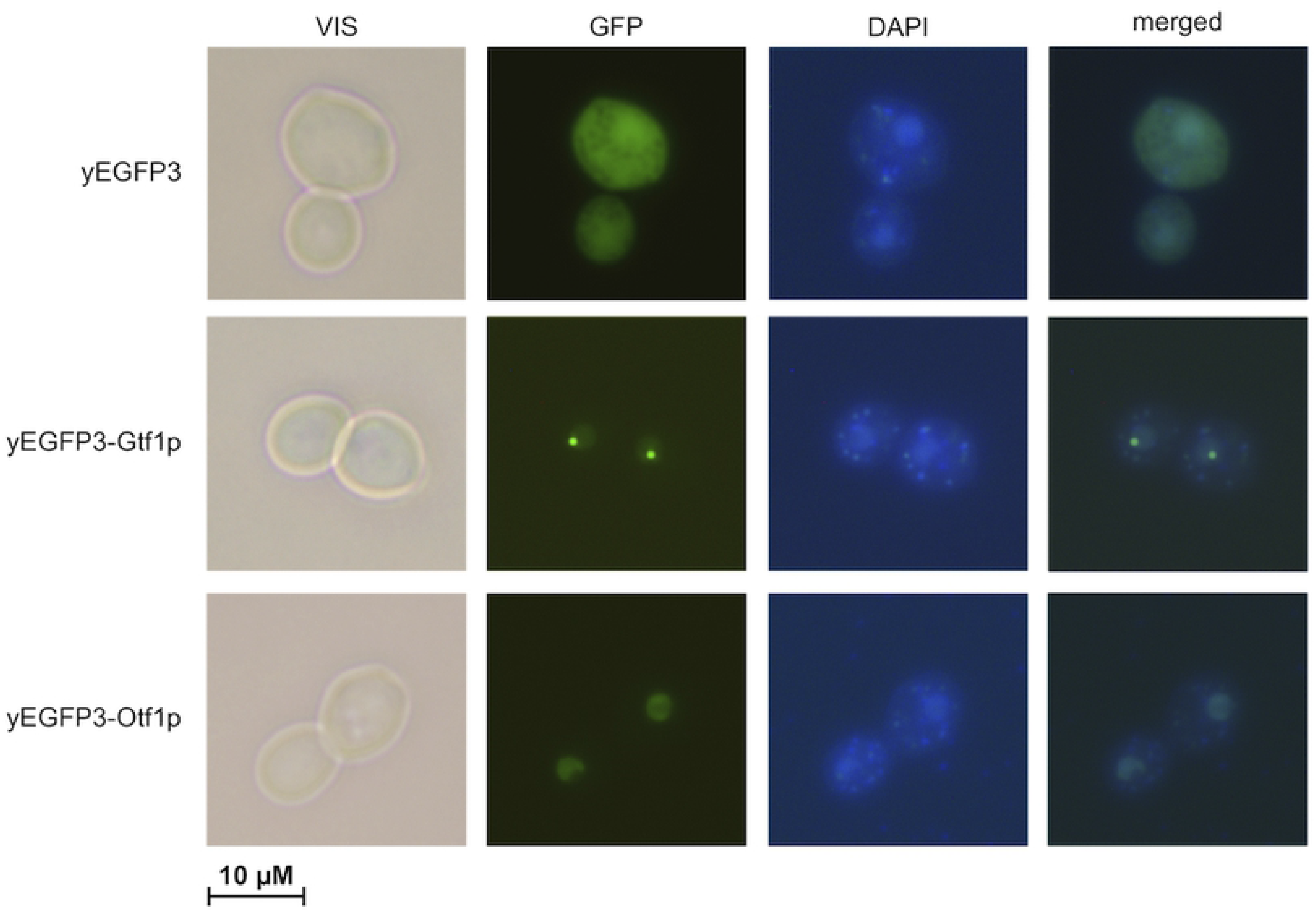
Transcription factors Otf1p and Gtf1 tagged with yEGFP3 at their N-termini localize in the cell nuclei. *C. parapsilosis* SR23 met1^-^ cells transformed with pPK6, pPK6-OTF1 or pPK6-GTF1 plasmids were grown overnight in an SGal medium. The cells were washed with water and DNA in cells was stained with DAPI.

To demonstrate that Otf1p and Gtf1p are involved in the transcriptional control of the 3-OAP and GP, respectively, we constructed knockout strains lacking both alleles of *OTF1* or *GTF1* and tested their ability to utilize different hydroxybenzenes and hydroxybenzoates as a sole carbon source. We found that the Δ*otf1*/Δ*otf1* mutant exhibits a growth defect on several substrates metabolized via the 3-OAP. While its growth is impaired in media containing hydroxybenzoates (i.e. 4-hydroxybenzoate, 2,4-dihydroxybenzoate, 3,4-dihydroxybenzoate), we did not observe a growth defect in media containing hydroxybenzenes (resorcinol, hydroquinone). On the other hand, the Δ*gtf1*/Δ*gtf1* mutant is unable to grow in media containing 3-hydroxybenzoate or gentisate, which are degraded via the GP (**Fig 8**). The phenotypes of both mutants indicate that Otf1p and Gtf1p are involved in the control of the 3-OAP and GP, respectively.

**Fig 8.**
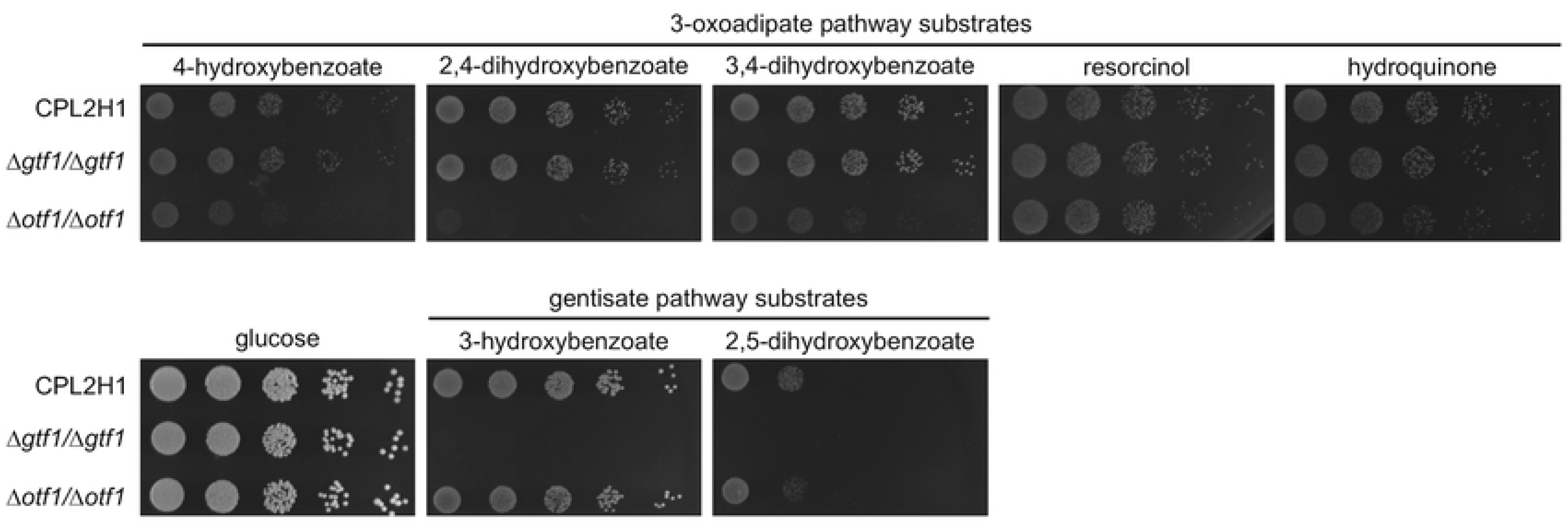
*C. parapsilosis* mutants Δ*otf1*/Δ*otf1* and Δ*gtf1*/Δ*gtf1* exhibit impaired growth on hydroxyaromatic substrates. Indicated strains were pre-grown overnight in a complex medium (YPD) at 28 °C, washed with water and resuspended to ∼6×10^6^ cells/ml. Serial fivefold dilutions were then spotted on solid synthetic media containing indicated carbon sources. The plates were incubated for 4 days at 28 °C.

To investigate the role of Otf1p and Gtf1p in the control of the 3-OAP and GP genes, respectively, we compared the transcriptomic profiles of the knockout mutants with the parental strain CPL2H1 cultivated in synthetic medium containing a mixture of hydroxyaromatic carbon sources (i.e. 3-hydroxybenzoate, 4-hydroxybenzoate, and hydroquinone) metabolized via the GP or 3-OAP. The RNA-Seq experiment demonstrated that expression of the genes present in the GP gene cluster is substantially decreased in the Δ*gtf1*/Δ*gtf1* mutant compared to the parental strain (**Fig 3, S2 Table**, **S6 Fig).** In addition, we found that the transcript(s) derived from *CANPARB_p44920-A* and *CANPARB_p44910-A* ORFs coding for a predicted amidohydrolase superfamily protein is almost absent in this mutant.

A comparison of the Δ*otf1*/Δ*otf1* mutant and CPL2H1 cells revealed more subtle differences in the expression of genes for the 3-OAP enzymes. We found that *MNX1* and *HBT2* are downregulated by 6.37- and 1.97-fold, respectively (**Fig 3**, **S2 Table**, **S6 Fig)**. As these genes code for 4-hydroxybenzoate 1-hydroxylase decarboxylating hydroxybenzoates to hydroxybenzenes (7,40,41) and a hydroxybenzoate transporter (10), their decreased expression goes in line with the observation that the Δ*otf1*/Δ*otf1* mutant has impaired growth on hydroxybenzoates (**Fig 7**). The expression of the genes *MNX3, HDX1*, *FRD1*, *OSC1*, and *OCT1* encoding remaining enzymes of the 3-OAP is also slightly decreased (i.e. by 1.45 to 1.90-fold). However, as the mutant grows on media with hydroquinone or resorcinol, we assume that expression of these genes is sufficient for utilization of both hydroxybenzenes.

### Otf1p and Gtf1p recognize specific motifs in promoter sequences

As described above, the genes coding for the enzymes of the 3-OAP and GP are highly upregulated in the cells assimilating hydroxyaromatic substrates. To identify potential regulatory motifs involved in their transcriptional control, we searched corresponding promoter sequences for putative Otf1p- and Gtf1p-binding sites. Both transcription factors belong to the Gal4-like family whose members recognize sequences containing CGG triplets oriented as inverted repeats separated by a distinct number of nucleotides, although other terminal nucleotides such as GGA were also identified in the binding sites (44, 45). Moreover, Otf1p is a homolog of the transcription factor *qa-1F* activating expression of the quinic acid gene cluster in *Neurospora crassa*, which recognizes a 16-mer motif GGATAATCGATTATCC (46). The search of the *MNX1* promoter sequence revealed a similar motif GGRN_10_WCC occurring at positions - 1376 to −1361 and −812 to −797. We also found that this motif is present in the promoters of other genes coding for the 3-OAP enzymes (i.e. *FRD1*, *HDX1, MNX3*, *OCT1, OSC1*), hydroxybenzoate transporter (*HBT2*) and its paralogs (*HBT3*, *HBT4*) (**S7 Fig**, **S8 Fig**) which are co-induced in the cells grown in media with 4-hydroxybenzoate or hydroquinone (**Fig 3**). Distribution of the GGRN_10_WCC motif indicates that it represents a potential binding site for the transcription factor Otf1p. To search for putative Gtf1p-binding sites, we analyzed the promoters of the GP cluster genes using the MEME tool (http://meme-suite.org; (47)) and identified a motif TCGGN_8_TCC (E-value = 2.7e-1) which occurs upstream of each ORF in the GP gene cluster, except for the *GTF1* gene. Alignment of the identified hits including their flanking sequences allowed us to refine the consensus sequence to a GGAN_7_TCC motif (**S7 Fig**, **S8 Fig**). Four putative Gtf1p-binding sites occur at positions −1111 to −1099, −428 to −416, −299 to −287, and −263 to −251 upstream of the *MNX2* coding sequence. As the first site is closer to the *HBT1* ORF (i.e. - 366 to −354), we assume that it may be involved in the control of this gene. Additional sites present in the GP gene cluster occur in the intergenic region between *GDX1* and *FPH1* (i.e. −191 to −179 and −201 to −189 upstream of *GDX1* and *FPH1* ORFs, respectively) and upstream of the *GFA1* ORF (i.e. −174 to −162 and −118 to −106). The motif is also present upstream of *CANPARB_p44920-A* (i.e. −184 to −172), which along with the GP cluster genes is also highly induced in media containing 3-hydroxybenzoate (**Fig 3**).

As our previous studies (7,9,10) indicated that the 3-OAP and GP genes are repressed in media containing glucose, we also searched the promoter sequences for sequence motifs potentially mediating this process. We have found several copies of the SYGGRG motif which is recognized by transcriptional repressors Mig1/Mig2 both in *S. cerevisiae* and *C. albicans* (48–51). Some of these sites (e.g. −139 to −134 and −108 to −103 upstream of *MNX2* and *HBT1* ORFs, respectively) are located near an A/T-box, which is known to be associated with bending of a DNA molecule upon Mig1/Mig2-binding (48) supporting the idea that at least some of the SYGGRG sites are functional in *C. parapsilosis*.

To demonstrate that transcription factors Otf1p and Gtf1p recognize the predicted motifs, we performed EMSA experiments using the protein extracts prepared from the wild type cells (CPL2H1) as well as the mutants lacking a functional copy of the corresponding transcription factor and the labeled ds-oligonucleotide probes OTF1-MNX1 and GTF1-MNX2 derived from the promoters of *MNX1* and *MNX2* genes, respectively. These probes contain a single copy of the predicted binding site. In DNA-binding reactions performed using the extract from the wild type cells we identified one and two bands using the probes OTF1-MNX1 and GTF1-MNX2, respectively (**Fig 9**). As these bands were absent when the extracts were prepared from the mutant cells (i.e. Δ*otf1*/Δ*otf1* for OTF1-MNX1; Δ*gtf1*/Δ*gtf1* for GTF1-MNX2), we assume that they correspond to the DNA-protein complexes containing the corresponding transcription factor and the probe. To further support this idea, we showed that the 50-fold and higher molar excess of unlabeled oligonucleotide used in the assay as a specific competitor outcompetes the labeled probe from the complex. Importantly, the oligonucleotides OTF1-MNX1_mut and GTF1_MNX2_mut carrying alterations in the conserved positions of predicted binding sites (i.e. TTTN_10_TAA and TTTN_7_AAA, respectively), did not interfere with the complex formation.

**Fig 9.**
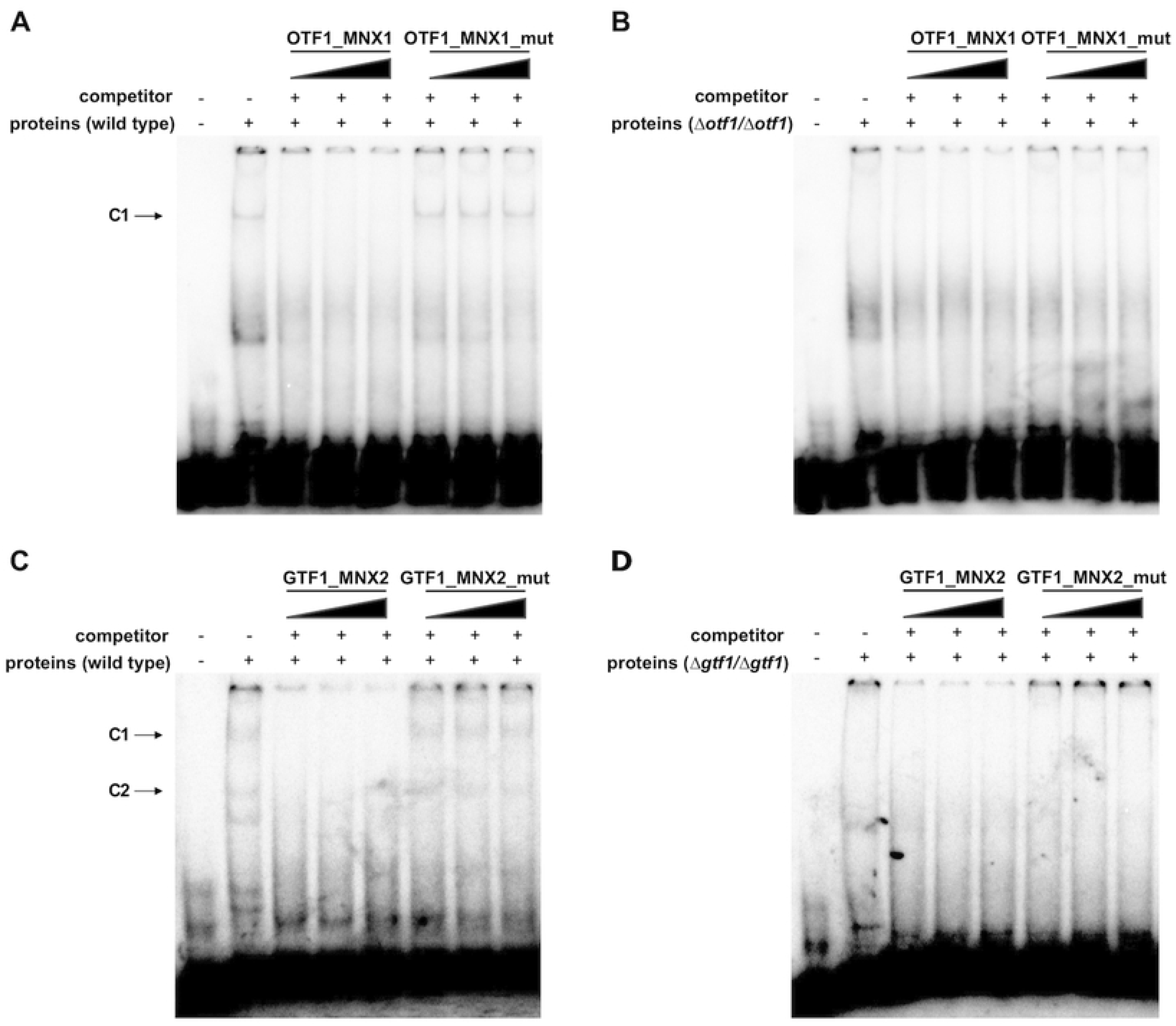
Transcription factors Otf1p and Gtf1p bind to predicted sequence motifs. The EMSA experiments were performed using the protein extracts prepared from CPL2H1 (A),(C), Δ*otf1*/Δ*otf1* (B), and Δ*gtf1*/Δ*gtf1* (D) cells and the 5’ end-labeled dsDNA probes containing the predicted Otf1p-binding site from the *MNX1* promoter (OTF1_MNX1; (A),(B)) or the Gtf1p-binding site from the *MNX2* promoter (GTF1_MNX2; (C),(D)). The ds oligonucleotide competitors containing either the wild type (OTF1_MNX1, GTF1_MNX2) or mutated binding motifs (OTF1_MNX1_mut, GTF1_MNX2_mut) were used with increasing amounts of 100, 300, and 500 ng as indicated above lanes.

Taken together, we demonstrate that Otf1p and Gtf1p are Gal4p-like transcription factors present in the extracts from the wild type cells and they specifically bind to DNA fragments carrying the motifs GGRN_10_WCC and GGAN_7_TCC, respectively. Gtf1p appears as the main transcriptional activator of the GP gene cluster. On the other hand, although Otf1p contributes to transcriptional activation of the 3-OAP genes, it predominantly controls the expression of *MNX1* encoding decarboxylating mononoxygenase (7, 41). This conclusion is supported by differences in the gene expression profiles of the cells grown on 4-hydroxybenzoate compared to those assimilating hydroquinone (**Fig 3**) and the growth phenotypes (**Fig 8**) and underscores the physiological differences in catabolic degradation of hydroxybenzenes and hydroxybenzoates. These results imply that, besides Otf1p, activation of the 3-OAP genes requires additional transcription factor(s). As 4-hydroxybenzoate and hydroquinone differ by the presence of carboxyl group, we speculate that bicarbonate anions generated upon Mnx1p-catalyzed decarboxylation of 4-hydroxybenzoate (**Fig 6B**) and corresponding cellular response may also contribute to identified differences.

### Phylogenetic analyses

The phylogenetic relationships of the transcription factors Otf1p and Gtf1p, as well as the twelve FAAs in *C. parapsilosis* were first assessed by investigating pre-computed phylogenies and orthology and paralogy relationships in MetaPhORs v2 (52) and PhylomeDB v4 (53) as of October 2020. As Otf1p and Gtf1p display a very sparse distribution among Saccharomycotina, we performed new phylogenetic reconstruction (see Materials and Methods) with the first 250 best Blast hits (e-value < 10^-20^) in a search against NCBI non-redundant database (as of October 2020). *GTF1* phylogeny (**Fig 10**) closely resembles that previously reported for other genes of the GP cluster such as *GDX1* (9), with a sparse distribution within Saccharomycotina and closely related to Pezizomycotina and Zygomycotina sequences, suggestive of a possible ancient horizontal gene transfer. *OTF1* phylogenetic reconstruction reveals a somewhat broader distribution within Saccharomycotina and a close relationship with Pezizomycotina sequences (**Fig 10**).

**Fig 10.**
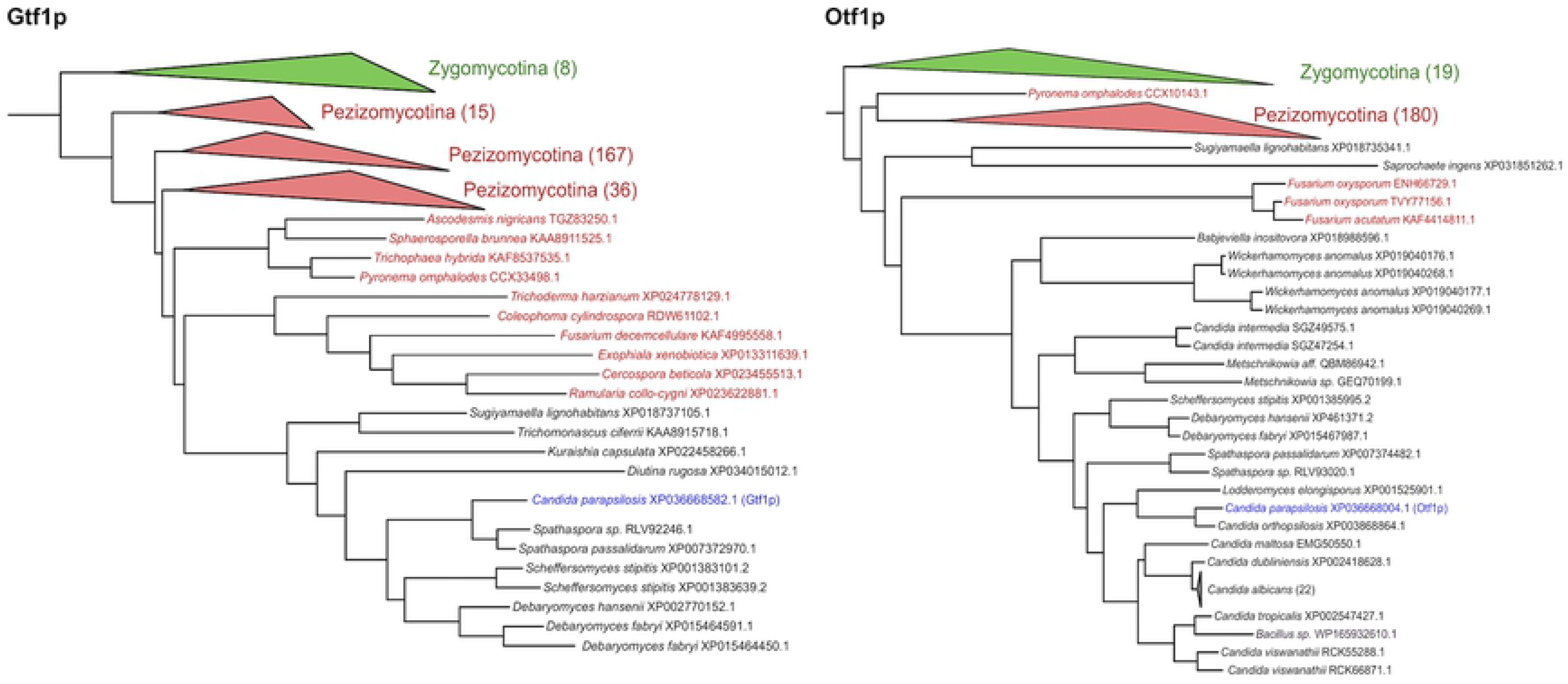
Phylogenetic relationships of the transcription factors Gtf1p (left), Otf1p (right), and their closest homologs. For simplicity, some monophyletic tree partitions including sequences from the same taxonomic classification are collapsed and their number is indicated in brackets. Zygomycotina sequences or partitions are indicated in green, Pezizomycotina sequences and partitions are indicated in red, Saccharomycotina sequences are indicated in black, and the *C. parapsilosis* sequence used as a seed in the blast searches is indicated in blue. Note several *C. albicans* sequences likely correspond to redundant sequences from different strains or sequencing projects. Note as well WP_165932610 sequence assigned to a *Bacillus* strain, which could correspond to a taxonomic miss-assignment or contamination. The full phylogenetic trees in newick format, including all sequence names and branch support is provided as supplemental information (**S1 Text**, **S2 Text**).

Previously we proposed that the 3-OAP variant occurring in *C. parapsilosis* emerged in an ancestral lineage before the divergence of the ‘CUG-Ser1’ clade from other Saccharomycotina lineages by an upgrade of a shorter version of this pathway (such as seen in *C. albicans*), which allows degradation of only hydroxybenzenes (10). The principal difference between the two variants is the presence of both the Mnx1p-catalyzed decarboxylation step and the functional uptake of hydroxybenzoates provided by Hbt2p and possibly also by its paralogs Hbt3p and Hbt4p, in the longer 3-OAP version. The acquisition of *OTF1* by horizontal gene transfer might have served the need for a specific regulation of this upgraded pathway. Differences in the transcriptional control of *MNX1* and to some extent also *HBT2* and *HBT3* compared to remaining 3-OAP genes (see above) supports the upgrade scenario and provides additional insight into the evolution of this pathway.

The evolutionary relationships of the twelve *FAA* genes in *C. parapsilosis* is well represented in PhylomeDB trees (see a simplified example in **S3 Fig**). This reveals an intricate evolution of this family with at least ten nested gene duplications at different ages leading to the twelve paralogs present in *C. parapsilosis* and with complex one-to-many orthology and paralogy relationships with the four *FAA* genes present in *S. cerevisiae* and *C. albicans*. This highlights a dynamic gene copy evolution leading to complexification of the FA metabolism in the *C. parapsilosis* clade.

Finally, we investigated the possible origin of the putative amidohydrolase gene (*CANPARB_p44920-A* and *CANPARB_p44910-A*) identified in this work. PhylomeDB searches rendered no results, but MetaPhOrs identified an ortholog in *C. metapsilosis* (g2237) sharing 64% protein identity. The *C. metapsilosis* gene has a single reading frame indicating that in *C. parapsilosis* ancestor the gene was split up into two ORFs by an in-frame stop codon UGA. In general, this alteration would inactivate a gene function, although stop codon bypassing or readthrough events (54) could generate a full-length protein corresponding to the polypeptide translated from uninterrupted ORF. Although both *C. parapsilosis* ORFs are transcribed on 3-hydroxybenzoate and the transcript is regulated by transcriptional activator Gtf1p, we did not identify peptides derived neither from individual ORFs nor from a deduced full-length protein by LC-MS/MS analysis. Searches in NCBI non-redundant database identified only bacterial sequences among the top 500 hits, with the best matches belonging to various *Pseudomonas* species with e-values ranging from 10^-105^ to 10^-102^ and sequence identities between 49 and 53% at the protein level. A multiple sequence alignment of the first 100 hits and the two *Candida* sequences revealed conservation of numerous amino acid residues (**S1 Fig**). This result indicates that this gene represents a relatively recent transfer of a gene encoding a putative amidohydrolase to the common ancestor of *C. parapsilosis* and *C. metapsilosis*. Alternatively, considering the relatively low sequence identity between the two transferred genes and their location in different, non-syntenic chromosomal locations in each of the species, two independent origins from a related bacterial donor might be postulated. The low levels of similarity exhibited between the transferred genes and their closest bacterial donors preclude us to pinpoint a specific bacterial species. Interestingly, when limiting the search to Saccharomycotina three significant hits appeared from sequences in the unrelated yeasts *Wickerhamiella sorbophila* (acc.no. XP_024665283.1)*, Trichomonascus ciferrii* (KAA8915622.1), and *Naumovozyma castellii* (XP_003673849.1). However, their sequence identities with the *C. parapsilosis* protein were lower than that of the bacterial homologs (39%, 35%, and 29%, respectively), suggesting they are more distantly related. Indeed, searches using these other yeast proteins as queries identified other bacteria (for *W. sorbophila*) or non-overlapping species of Pezizomycotina fungi (for *T. ciferrii* and *N. castellii*) among the first 100 significant hits, suggesting each of these yeasts acquired a different amidohydrolase gene in independent horizontal gene transfers. Such recurrent horizontal gene transfer scenario is reminiscent of other metabolic genes including amino acid racemases, which are also present in *C. parapsilosis* and other yeast species (55).

## Materials and Methods

### Yeast strains

*C. parapsilosis* strains CLIB214 (identical to the type strain CBS604), CPL2H1 (CLIB214 Δ*leu2*/Δ*leu2*, Δ*his1*/Δ*his1*; (56)), its mutant derivatives (Δ*gtf1*/Δ*gtf1* and Δ*otf1*/Δ*otf1*; this study), CDU1 (CLIB214 Δ*ura3*/Δ*ura3*; (57)), and SR23 met1^-^ (*ade^-^, lys4^-^, met1^-^*; (58)) were used in this study.

### DNA isolation and sequencing

Genomic DNA was isolated from the strain CLIB214. Briefly, a yeast culture grown overnight at 28 °C in 100 ml of YPD medium (1% [wt/vol] yeast extract, 2% [wt/vol] peptone, 2% [wt/vol] glucose) at 28 °C was harvested by centrifugation (5 min, 2,100 *g* at 4 °C), the cells were resuspended in 20 ml of 2% [vol/vol] 2-mercaptoethanol and incubated for 30 min at room temperature. The spheroplasts were prepared in 6 ml of 1 M sorbitol, 10 mM EDTA (pH 8.0) containing 0.125 mg of Zymolyase 20T (Seikagaku) at 37 °C, pelleted by centrifugation (5 min, 2,100 *g* at 4 °C) and lysed in 3 ml of 0.15 M NaCl, 0.1 M EDTA (pH 8.0), 0.1% [wt/vol] SDS. Proteins were removed by three extractions with equal volume of phenol buffered with 10 mM Tris-HCl, 1 mM EDTA (pH 8.0) and by one extraction with equal volume of chloroform : isoamyl alcohol (24 : 1). Nucleic acids were precipitated using 0.1 M NaCl and 1 volume of 96% [vol/vol] ethanol, pelleted by centrifugation (10 min, 16,100 *g* at 4 °C), washed with 70% [vol/vol] ethanol and air dried. The precipitate was dissolved in 1 ml of 10 mM Tris-HCl, 1 mM EDTA (pH 7.5), 0.1 mg/ml RNase A and incubated for 45 min at 37 °C. DNA was extracted by phenol and chloroform : isoamyl alcohol, precipitated using 0.1 M NaCl and 2 volumes of 96% [vol/vol] ethanol, washed with 70% [vol/vol] ethanol, air dried, dissolved in 150 μl 10 mM Tris-HCl, 1 mM EDTA (pH 7.5) and purified on a Genomic-tip 100/G (Qiagen) according to the manufacturer’s instructions. A paired-end (2×151-nt) TruSeq PCR-free DNA library was sequenced on a NovaSeq6000 platform at Macrogen Korea, yielding 81,578,508 reads (12.32 Gbp; 944x coverage). Nanopore sequencing was performed on a MinION Mk-1B device with an R9.4.1 flow cell using a Rapid barcoding kit (SQK-RBK004; Oxford Nanopore Technologies). 119,788 reads were obtained (mean and median lengths are 9,200.2 and 5,938 nucleotides, respectively) totaling 1.1 GBp (84x coverage). Nanopore reads were assembled by Canu version 1.9 (59), resulting in 20 contigs, which were manually examined. Chromosomes 1, 2, 3, 4 and 7 were used as assembled by Canu. Chromosome 8 was created by connecting two Canu contigs. In the contig corresponding to chromosome 5, a 8 kbp region was replaced by a longer 14.5 kb version from one of the excluded shorter contigs. This region contains two copies of the *PDR5* gene, possibly with a copy number variation. Finally, regions directly upstream and downstream of ribosomal DNA (rDNA) arrays on chromosome 6 were misassembled in the Canu assembly. These regions were replaced by sequences assembled from Illumina reads by SPAdes version 3.12 (60). After these manual modifications, the entire assembly was polished first by Medaka version 0.11.5 (61) using nanopore reads, and then by three iterations of Pilon version 1.12 (62). The rDNA repeat poses problems for polishing due to its repetitive nature, and thus a single copy of the repeat was polished separately by Pilon and then used in the final assembly. Mitochondrial DNA was taken from the GenBank acc. no. DQ376035.2. A whole genome alignment to the reference genome sequence from the strain CDC317 (GCA_000182765.2; (21, 63)) was computed by Last version 830 (64) followed by last-split to assign to each portion of CDC317 sequence a unique position in our assembly. To annotate protein coding genes, the genes from the strain CDC317 were aligned to our assembly by Blat version v. 36×2 (65) and supplied as hints to Augustus version 3.2.3 (66). Augustus was run originally with parameters for *Candida albicans*, then retrained on the predictions matching CDC317 genes. Disagreements with CDC317 annotation were manually inspected, and as a result, 61 genes were modified, 12 genes removed and 72 genes added.

### Electrophoretic karyotyping

About 1×10^9^ cells of the strain CLIB214 grown overnight in a YPD medium were resuspended in 0.5 ml of 1.2 M sorbitol, 40 mM citric acid, 120 mM disodium phosphate, 20 mM EDTA (pH 8.0), 5 mM DTT, 0.2 mg/ml zymolyase 20T (Seikagaku) and incubated for 90 min at 37 °C. Protoplasts were then harvested by centrifugation (1 min, 2,100 *g*), resuspended in 1 ml of molten 1% [wt/vol] low melting point agarose in 50 mM EDTA (pH 8.5) cooled to 45 °C and poured into the sample forms. The agarose embedded samples were incubated for 30 min at 37 °C in 10 mM Tris.Cl, 0.5 M EDTA (pH 8.5) and then overnight at 50 °C in 1% [wt/vol] N-lauroylsarcosine, 0.5 M EDTA (pH 8.5), 0.5 mg/ml proteinase K. Pulsed-field gel electrophoresis (PFGE) was performed in a CHEF Mapper XA Chiller System (Bio-Rad) with 120° angle between the electric fields at the following settings: (I) 120 s pulses for 24 h followed by 240 s pulses for 36 h at 4.5 V/cm; (II) 120 s pulses for 20 h followed by 240 s pulses for 28 h at 4 V/cm; (III): 60 s pulses for 15 hours followed by 90 s pulses for 9 hours at 6 V/cm. The samples were separated in 0.8% (settings I and II) or 1% [wt/vol] agarose gels (settings III) in 0.5x TBE buffer (45 mM Tris-borate, 1 mM EDTA, pH 8.0) at 14 °C, throughout. After PFGE, the gels were stained with ethidium bromide (0.5 μg/ml) for 45 min and washed in water for additional 45 min.

### RNA-Seq analysis

*C. parapsilosis* CLIB214 was pre-cultivated in an SD medium (0.67% [wt/vol] yeast nitrogen base w/o amino acids (Difco), 2% [wt/vol] glucose) for 24 h at 28 °C, then washed in water and re-grown in SMix10 medium (0.67% [wt/vol] yeast nitrogen base w/o amino acids (Difco), 3.3 mM 3-hydroxybenzoate, 3.3 mM 4-hydroxybenzoate, and 3.3 mM hydroquinone) for additional 24 h at 28 °C. The pre-culture was inoculated (OD_600_ ∼ 0.3) in triplicates into synthetic media (0.67% [wt/vol] yeast nitrogen base w/o amino acids (Difco)) containing 2% [wt/vol] galactose (SGal) or 10 mM hydroxyaromatic compound (i.e. 3-hydroxybenzoate (S3OH), 4-hydroxybenzoate (S4OH) or hydroquinone (SHyd)) as a sole carbon source and cultivated at 28 °C till OD_600_ ∼ 1. The consumption of hydroxyaromatic compounds in the cultivation media was analyzed spectrophotometrically (**S9 Fig**). The cultures of CPL2H1, Δ*gtf1*/Δ*gtf1,* and Δ*otf1*/Δ*otf1* were prepared similarly, except that the cultivation media were supplemented with leucine (20 μg/ml) and histidine (20 μg/ml) and the cells were grown in SMix15 medium (0.67% [wt/vol] yeast nitrogen base w/o amino acids (Difco), 5 mM 3-hydroxybenzoate, 5 mM 4-hydroxybenzoate, and 5 mM hydroquinone). Total RNA was isolated by extraction with hot acid phenol essentially as described in (67) and purified using an RNeasy mini kit (Qiagen) according to the manufacturer’s instructions. Transcriptome sequencing reads were generated from TruSeq stranded mRNA LT paired-end (2×151-nt) libraries on a NovaSeq6000 platform at Macrogen Korea. Reads were processed with Trimmomatic 0.39 (68) and mapped to the CLIB214 genome sequence using HiSat2 2.1.0 (69). Duplicated reads were removed using samtools rmdup and the coverage was calculated using samtools depth (samtools version 1.9; (70)). Differential gene expression analysis was performed using the Geneious 11.1.5 package (Biomatters); the DESeq2 method (71) was used for samples in biological triplicates (*i.e.* CLIB214 grown in SGal, S3OH, S4OH, and SHyd media), the Geneious method was used for comparisons of CPL2H1 vs. Δ*gtf1*/Δ*gtf1* and CPL2H1 vs. Δ*otf1*/Δ*otf1* grown in SMix15 medium. Heatmaps were generated using the pheatmap package 1.0.12 (https://CRAN.R-project.org/package=pheatmap; (72)). Gene ontology and KEGG database searches were performed using the Candida Genome Database GO Slim Mapper and Term Finder (http://www.candidagenome.org) and the KEGG mapper (https://www.genome.jp/kegg/tool/map_pathway.html), respectively.

### Analyses of cultivation media

The consumption of hydroxyaromatic compounds and pH in cultivation media were analyzed before and at the end of cultivation using a Multiskan GO spectrophotometer (Thermo Scientific) and PH CHECK pH meter (Dostmann Electronic), respectively. The measurements were performed at room temperature. Following absorption maxima and media dilutions were used in the substrate consumption analyses; A_max_=297 nm, 2-fold dilution (3-hydroxybenzoate), A_max_=255 nm, 20-fold dilution (4-hydroxybenzoate), and A_max_=290 nm, 5-fold dilution (hydroquinone). To monitor pH changes during cultivations, synthetic media (see above) varying by the carbon source were supplemented with 0.01% [wt/vol] bromothymol blue and adjusted to pH 6.1-6.4 with NaOH. The cultures were inoculated to 6×10^6^ cells/ml and grown at 28 °C to mid exponential phase. Cell-free media were used as a control. To document color, the cultures were centrifuged (1 min, 2,100 *g*) to remove the cells and 100 µl of cultivation media were transferred into wells of a 96-well plate and photographed using a Nikon D7000 camera.

### Proteomic analysis

Protein extracts were prepared in triplicates from *C. parapsilosis* CLIB214 cells pre-cultivated overnight at 28 °C in an S3OH medium, inoculated (5×10^6^ cells/ml) to SGal, S3OH, S4OH, and SHyd media and grown at 28 °C till ∼ 10^7^ cells/ml. The cells were harvested by centrifugation (5 min, 2,100 *g* at 4 °C), resuspended in 50 mM Tris-HCl (pH 8.8), 1 mM EDTA (pH 8.0) and homogenised using FastPrep-24 (MP Biomedicals). Cell debris was removed by centrifugation (15 min, 16,000 *g* at 4 °C) and protein concentration was determined using Bradford’s method (73). For LC-MS/MS analysis, protein aliquots (50 µg) were diluted in 100 µl of 25 mM Tris-HCl (pH 7.8), 0.1 mM CaCl_2_, treated using 5 mM dithiothreitol for 30 min at 60 °C and alkylated in 40 mM chloroacetamide for 1 hour at 37 °C. The proteins were digested overnight by trypsin (1:30 [wt/wt]) at 37 °C. Acidified (0.5% [vol/vol] trifluoroacetic acid (TFA)) peptide solution was clarified by centrifugation and purified on a microtip C18 SPE. The concentration of eluted peptides was determined by Pierce™ Quantitative Fluorometric Peptide Assay (Thermo Scientific). The peptides were dissolved in 0.1% [vol/vol] TFA and 2% [vol/vol] acetonitrile (ACN), loaded (500 ng per run) onto a trap column (PepMap100 C18, 300 μm x 5 mm, Dionex, CA, USA) and separated with an EASY-Spray C18 column (75 µm x 500 mm, Thermo Scientific) on Ultimate 3000 RSLCnano system (Dionex) in a 120-minute gradient (3-43% B), curve 7, and flow-rate 250 nl/min. The two mobile phases were used: 0.1% [vol/vol] formic acid (A) and 80% [vol/vol] ACN with 0.1% [vol/vol] formic acid (B). Eluted peptides were sprayed directly into Orbitrap Elite mass spectrometer (Thermo Scientific, MA, USA) and spectral datasets were collected in the data dependent mode using Top15 strategy for the selection of precursor ions for the HCD fragmentation (74). Each of the three experimental replicates was analysed in technical triplicates. Protein spectra were analyzed by MaxQuant software (version 1.6.17.0) using carbamidomethylation (C) as permanent and oxidation (M) and N-terminal acetylation as variable modifications, with engaged ‘match between the runs’ feature and label-free quantification (LFQ) and further examined in Perseus version 1.6.15.0 (75, 76). The search was performed against the *C. parapsilosis* CLIB214 protein database containing 5856 entries. Proteins were evaluated and annotated based on information from CDC317 strain orthologs. Contaminating peptides, reverse peptides and peptides only identified by site were removed, then the protein entries were further filtered to have at least two LFQ values in at least one of the biological conditions (different carbon sources). Following an imputation, differentially expressed proteins were identified by ANOVA test (permutation-based FDR 0.01).

### Preparation of knockout strains

The mutants lacking either *GTF1* or *OTF1* gene were generated in the strain CPL2H1 essentially as described in (56, 77). Deletion constructions contained the upstream (UpFw primer and UpRev primer; **S4 Table**) and downstream (DownFw primer and DownRev primer) homologous regions of the target ORF and either *Candida dubliniensis HIS1* or *Candida maltosa LEU2* sequences as selection markers. For selection marker amplification the primers ‘pSN52/pSN40 Fw’ and ‘pSN52/pSN40 Rev’ were used. DownFw, UpRev, and the primers used for marker amplification also harbored fusion sequences for later fragment joining. The reverse primer (’pSN52/pSN40 Rev’) used for marker amplification also carried a TAG sequence between the mentioned fusion sequences. Deletion cassettes were transformed into CPL2H1 strain and the transformants were plated onto selective media. Heterozygous mutants were obtained and used to prepare homozygous mutants. Mutant strains were verified by colony polymerase chain reaction (PCR) using the primers specific for both the marker sequences and the outside of the integration sites at both the upstream and downstream homologous regions. The ORF specific primer ‘5’-check primer’ was used as forward primer together with ‘*LEU1/HIS1 primer’* as reverse primer, while the ORF specific primer ‘3’-check primer’ was applied as reverse primer together with the ‘*LEU2/HIS2* primer’ as forward primer.

Assimilation tests of the wild type and mutant strains were performed on solid synthetic media (0.67% [wt/vol] yeast nitrogen base w/o amino acids (Difco), 2% [wt/vol] agar, 30 μg/ml leucine, 20 μg/ml histidine) differing by the carbon source (i.e. 2% [wt/vol] glucose (SD), 10 mM 3-hydroxybenzoate (S3OH), 10 mM 4-hydroxybenzoate (S4OH), 10 mM 2,4-dihydroxybenzoate (S24diOH), 10 mM 2,5-dihydroxybenzoate (S25diOH), 10 mM 3,4-dihydroxybenzoate (S34diOH), 10 mM hydroquinone (SHyd) or 10 mM resorcinol (SRes)). Prior to the addition to the media, hydroxyaromatic compounds were dissolved in dimethyl sulfoxide (DMSO) as 0.5 M stocks.

### Fluorescence microscopy

The cells were observed using a BX50 microscope with the appropriate filter set and a digital camera DP70 (Olympus Optical). To visualize peroxisomes in *C. parapsilosis* cells, we constructed a plasmid pBP7-mCherry-SKL expressing the mCherry protein tagged with peroxisomal targeting signal ‘SKL’ at its C-terminus and a control plasmid pBP7-mCherry expressing the unmodified protein. The mCherry coding sequence was amplified by PCR using the primers shown in **S4 Table** and the plasmid pMG2254 (39) as a template. The PCR products were inserted into the *Xba*I site of the pBP7 vector (78) using a Gibson assembly cloning kit (New England Biolabs). The cloned genes are placed downstream of the *GAL1* promoter in the resulting plasmid constructs. The constructs were transformed into *C. parapsilosis* cells CDU1 by the standard protocol (79). The transformants were cultivated overnight in liquid synthetic medium (0.67% [wt/vol] yeast nitrogen base w/o amino acids (Difco)) containing 2% [wt/vol] glucose (SD) at 28 °C. The cells were then inoculated to synthetic media differing by the carbon source (i.e. 2% [wt/vol] galactose (SGal), 10 mM 3-hydroxybenzoate (S3OH), 10 mM 4-hydroxybenzoate (S4OH)), cultivated for 24 hour (SGal), 48 hours (S3OH) or 72 hours (S4OH) at 28 °C and examined by fluorescence microscopy. To investigate the intracellular localization of Gtf1p and Otf1p, we constructed yEGFP3-tagged versions of both proteins as follows. The coding sequences of *GTF1* and *OTF1* were PCR-amplified from the CLIB214 genomic DNA using gene specific primers (**S4 Table**) and the PCR products were inserted into the *Sma*I site of the pPK6 vector (78) using a Gibson assembly cloning kit (New England Biolabs). This allows the expression of cloned genes under the control of the *GAL1* promoter. The plasmid constructs were transformed into *C. parapsilosis* cells SR23 met1^-^ as described in (79). The transformants were grown overnight in SGal medium (0.67% [wt/vol] yeast nitrogen base w/o amino acids (Difco), 1% [wt/vol] galactose) at 28 °C. Prior to fluorescent microscopy, the cellular DNA was stained with 4′,6-diamidino-2-phenylindole (DAPI, 1 µg/ml) for 20 min.

### Electrophoretic Mobility Shift Assay (EMSA)

The wild type (CPL2H1) and mutant (Δ*gtf1/Δgtf1*, Δ*otf1/Δotf1*) cells were grown in synthetic media containing combinations of hydroxyaromatic substrates (i.e. 7.5 mM 3-hydroxybenzoate and 7.5 mM 4-hydroxybenzoate (wild type); 2.5 mM 3-hydroxybenzoate, 2.5 mM 4-hydroxybenzoate, and 10 mM hydroquinone (Δ*gtf1/Δgtf1*); 10 mM 3-hydroxybenzoate and 5 mM 4-hydroxybenzoate (Δ*otf1/Δotf1*)) supplemented with leucine (40 μg/ml) and histidine (40 μg/ml). Protein extracts were prepared according to Winkler *et al.* (80) with some modifications. Ice-cold solutions were used throughout the experiment and all incubations were performed on ice. Cells were harvested at exponential growth phase by centrifugation (10 min, 3,600 *g* at 4 °C), washed with water, resuspended in 5 volumes of 200 mM Tris-HCl (pH 8.0), 400 mM (NH_4_)_2_SO_4_, 10 mM MgCl_2_, 1 mM EDTA, 7 mM 2-mercaptoethanoI, 10% [vol/vol] glycerol, 1 mM phenylmethylsulfonyl fluoride (PMSF), 1 × cOmplete^TM^ protease inhibitor cocktail tablet (Roche Applied Science). The cells were disrupted by vortexing with glass-beads (0.45-0.5 mm in diameter, 0.8 g/ml) 7 times for 1 min with intermittent cooling on ice for 1 min. Lysates were incubated for 30 min, centrifuged at 9,000 *g* for 60 min and proteins in supernatant were precipitated by addition of (NH_4_)_2_SO_4_ in 10 mM HEPES (pH 8.0), 5 mM EDTA, 1 mM PMSF for 30 min (the final concentration of (NH_4_)_2_SO_4_ was 40% [wt/vol] in total volume of 1.5 ml). The sample was centrifuged at 9,000 *g* for 15 min and the pellet was resuspended in 100 – 150 µl of 10 mM HEPES (pH 8.0), 5 mM EDTA, 7 mM 2-mercaptoethanoI, 20% [vol/vol] glycerol, 1 mM PMSF, 1× cOmplete^TM^ protease inhibitor cocktail tablet (Roche Applied Science). The protein extracts were stored at −80 °C prior to the use in DNA-binding assays. Oligonucleotide probes were prepared as follows. Direct strand oligonucleotides (**S4 Table**) were labeled at 5′ end by T4 polynucleotide kinase (Thermo Scientific) and [γ-^32^P]ATP (Hartmann Analytic), mixed with 3-fold molar excess of the unlabeled complementary oligonucleotide, heated at 100 °C for 10 min and slowly cooled down to room temperature to allow efficient formation of the double-stranded probes. The probes were purified using Illustra™ MicroSpin™ G-25 Columns (GE Healthcare). The DNA binding assays were carried out in 10 µl of 10 mM Tris-HCl (pH 7.5), 50 mM NaCl, 0.1 mM EDTA containing 15 µg of proteins, 2 ng of the ^32^P-labeled probe, 2 µg of poly(dA-dC) • poly(dG-dT). Unlabeled double-stranded oligonucleotides were used as specific competitors. The reaction mixtures were incubated for 15 min at room temperature and immediately loaded on 5% polyacrylamide gels in TG buffer (25 mM Tris-HCl (pH 8.3), 192 mM glycine). The electrophoresis was performed at 4 °C in the TG buffer at 10 V/cm for 90 min. Gels were fixed with 10 ml of 10% [vol/vol] methanol, 10% [vol/vol] acetic acid for 10 min, dried and exposed to the storage phosphor screen. Signal was detected using a Personal Molecular Imager FX (Bio-Rad).

### Phylogenetic analysis

Sequences were aligned with Muscle v3.8 (81) with default parameters and maximum likelihood phylogenetic trees were built using Iqtree v2.0 (82) allowing full exploration of model parameters and estimating the support of tree partitions using ultrafast bootstrap support with 1000 iterations (83). Orthology and paralogy relationships, as well as duplication nodes were inferred with the species overlap algorithm (84), with the relative age inferred from topological analysis (85). Blast searches were performed at NCBI website (https://blast.ncbi.nlm.nih.gov/) using default parameters unless indicated otherwise.

### Data availability

The CLIB214 genome assembly, nanopore and Illumina reads were deposited in the European Nucleotide Archive (ENA) under the project PRJEB37287. RNA-Seq data were submitted to ArrayExpress under the accessions E-MTAB-9442 and E-MTAB-9443. The mass spectrometry proteomics data have been deposited to the ProteomeXchange Consortium via the PRIDE (86) partner repository with the dataset identifier PXD024608 and 10.6019/PXD024608.

## Supporting information captions

**S1 Table. RNA-Seq analysis of *C. parapsilosis* cells.**

Lists of genes differentially expressed in *C. parapsilosis* CLIB214 cells grown in synthetic media containing 3-hydroxybenzoate, 4-hydroxybenzoate or hydroquinone compared to the cells assimilating galactose (i.e. S3OH vs. SGal, S4OH vs. SGal, SHyd vs. SGal) or 4-hydroxybenzoate (S3OH vs. S4OH, SHyd vs. S4OH). Note that only the genes exhibiting statistically significant log_2_ fold change values (p ≤ 0.05) are shown.

**S2 Table. RNA-Seq analysis of *C. parapsilosis* mutants lacking Otf1p or Gtf1p.**

Lists of genes differentially expressed in the *C. parapsilosis* mutants Δ*gtf1/Δgtf1* and Δ*otf1/Δotf1* compared to the parental strain CPL2H1 (i.e. Δ*gtf1/Δgtf1* vs. CPL2H1 and Δ*otf1/Δotf1* vs. CPL2H1). The cells were grown in synthetic media (SMix15) containing three hydroxyaromatic carbon sources (i.e. 3-hydroxybenzoate, 4-hydroxybenzoate, and hydroquinone). Note that only the genes exhibiting statistically significant log_2_ fold change values (p ≤ 0.05) are shown.

**S3 Table. LC-MS/MS analysis of proteins extracted from *C. parapsilosis* cells.**

Lists of proteins identified in the extracts of *C. parapsilosis* CLIB214 cells grown in synthetic media containing 3-hydroxybenzoate (S3OH), 4-hydroxybenzoate (S4OH), hydroquinone (SHyd) or galactose (SGal) as a carbon source. Protein spectra were subjected to label-free quantification (LFQ) and statistically evaluated.

**S4 Table. List of synthetic oligonucleotides.**

**S1 Text. Phylogenetic tree of GTF1 sequences in the Newick format.**

**S2 Text. Phylogenetic tree of OTF1 sequences in the Newick format.**

**S1 Fig. Homologs of amidohydrolase family proteins.**

Amino acid sequence alignment of conceptual translation of *C. parapsilosis* CANPARB_p44920-A (red shading), short intergenic spacer, and CANPARB_p44910-A (blue shading) with yeast (*C. metapsilosis* (g2237), *T. ciferrii* (KAA8915622.1), *W. sorbophila* (XP_024665283.1)*, N. castellii* (XP_003673849.1)) and bacterial (*Pseudomonas aestus* (P308_18355), *Paraburkholderia megapolitana* (SAMN05192543_101920), and *Variovorax* sp. (VAR608DRAFT_1163)) homologs. The alignment was calculated using MAFFT v7.450 (87).

**S2 Fig. Expression profiles of *C. parapsilosis* genes coding for metabolic enzymes.**

The heatmaps show the expression profiles obtained by the RNA-Seq and LC-MS/MS analyses. The log_2_ fold change values obtained by the RNA-Seq analysis (**S1 Table**) are shown on the left panel. Only the genes that are upregulated (log_2_ fold change ≥ 2; p ≤ 0.05) on at least one hydroxyaromatic substrate and code for protein products classified as metabolic enzymes (based on the searches using the BlastKOALA (https://www.kegg.jp/blastkoala/; (88)) and KEGG Mapper tools (https://www.kegg.jp/kegg/tool/map_pathway.html; (89)) are included. Note that the values that are not statistically significant (i.e. p > 0.05) are shown in parentheses. The values on the right panel represent log_2_ of mean LFQ intensity ratios taken from the LC-MS/MS analysis (**S3 Table**). Note that the LFQ values imputed from a normal distribution were used for proteins that were not identified on all carbon sources (shown in parentheses). Proteins CANPARB_p24940-A and CANPARB_p24960-A, and similarly also CANPARB_p56420-A and CANPARB_p56500-A, have almost identical sequences and therefore could not be distinguished by the LC-MS/MS analysis. Orthologs or best hits (indicated by an asterisk) from the *C. parapsilosis* reference strain CDC317, *C. albicans*, and *S. cerevisiae*, and the KEGG IDs are indicated.

**S3 Fig. Acyl-CoA synthetases in *C. parapsilosis*.**

(A) The heatmaps show the expression profiles of *C. parapsilosis FAA* genes. The log_2_ fold change values obtained by the RNA-Seq analysis (**S1 Table**) are shown on the left panel. Note that the values that are not statistically significant (i.e. p > 0.05) are shown in parentheses. The values on the right panel represent log_2_ of mean LFQ intensity ratios taken from the LC-MS/MS analysis (**S3 Table**). (B) Phylogenetic relationships of *C. parapsilosis FAA* genes and their homologs in other yeasts. The *CPAR2_200640* gene tree in phylome 498 from PhylomeDB (*Candida inconspicua* genome, described in (90)) was used as a template to create this figure, which is only shown partially here. Sequences from *C. parapsilosis* (black), *C. albicans* (red), and *S. cerevisiae* (blue) are highlighted with their names. Shadowed rectangles around them indicate, respectively, the spread of species from the *C. parapsilosis sensu lato*, *C. albicans / C. dubliniensis / C. tropicalis* clade, and *Saccharomyces / Nakaseomyces* clade. Colored circles indicate duplication nodes, with different colors indicating the relative age inferred from this duplication (see legend).

**S4 Fig. Expression profiles of *C. parapsilosis* genes involved in the biogenesis and metabolism of peroxisomes.**

The heatmaps show the expression profiles obtained from the RNA-Seq and LC-MS/MS analyses. The log_2_ fold change values obtained by the RNA-Seq analysis (**S1 Table**) are shown on the left panel. Only the genes that are upregulated (log_2_ fold change ≥ 2; p ≤ 0.05) on at least one hydroxyaromatic substrate and code for protein products classified into categories ‘peroxisome’, ‘peroxisomal matrix’, ‘peroxisomal membrane’ or ‘peroxisomal importomer complex’ (based on the GO subcellular component analysis; http://www.candidagenome.org) are included. Note that the values that are not statistically significant (i.e. p > 0.05) are shown in parentheses. The values on the right panel represent log_2_ of mean LFQ intensity ratios taken from the LC-MS/MS analysis (**S3 Table**). Orthologs or best hits (indicated by an asterisk) from the *C. parapsilosis* reference strain CDC317, *C. albicans*, and *S. cerevisiae* are shown.

**S5 Fig. Amino acid sequence alignments of Otf1p and Gtf1p orthologs.**

(A) Amino acid sequence alignment of C. parapsilosis Otf1p with the counterparts from C. metapsilosis (CMET_1974), C. orthopsilosis (CORT0C05870), C. albicans (ZCF10), C. tropicalis (CTRG_01734), Scheffersomyces stipitis (PICST_62477), and Spathaspora passalidarum (SPAPADRAFT_137814).

(B) Amino acid sequence alignment of C. parapsilosis Gtf1p with the counterparts from C. metapsilosis (CMET_1081), S. passalidarum (SPAPADRAFT_53773), Debaryomyces hansenii (DEHA2C00946g), and S. stipitis (PICST_57167 and PICST_65252). The alignments were calculated using MAFFT v7.450 (87). The GAL4-like domain (red shading) and fungal specific transcription factor domain (blue shading) were predicted using SMART 8.0 (91). Nuclear localisation signal (NLS, shown in magenta) was identified using SeqNLS (92).

**S6 Fig. Downregulated genes in *C. parapsilosis* mutants lacking Otf1p or Gtf1p.**

The heatmap shows the genes downregulated (log_2_ fold change ≤ −2; p ≤ 0.05; **S2 Table**) in the mutantsΔ*gtf1/Δgtf1* and Δ*otf1/Δotf1* compared to the parental strain CPL2H1 (Δ*gtf1/Δgtf1* vs. CPL2H1 and Δ*otf1/Δotf1* vs. CPL2H1). The cells were grown in an SMix15 medium containing three hydroxyaromatic carbon sources (i.e. 3-hydroxybenzoate, 4-hydroxybenzoate, and hydroquinone). Note that the values that are not statistically significant (i.e. p > 0.05) are shown in parentheses. Orthologs or best hits (indicated by an asterisk) from *C. parapsilosis* CDC317, *C. albicans,* and *S. cerevisiae,* and KEGG IDs are indicated.

**S7 Fig. Putative binding sites for Otf1p and Gtf1p in the promoters of 3-OAP and GP genes.** The occurrence of putative Otf1p (A; shown as blue triangles) and Gtf1p (B; blue rectangles) binding sites in the upstream regions of the genes encoding the components of the 3-OAP and GP, respectively. Putative Mig1p-binding sites (red triangles) and the positions of probes used in the EMSA experiments (black rectangles) are also depicted.

**S8 Fig. Predicted Otf1p and Gtf1p binding motifs.**

(A) Otf1p binding motif. (B) Gtf1p binding motif. The sequence logos were derived from predicted binding sites identified in the promoter sequences shown in **S7 Fig**.

**S9 Fig. Consumption of hydroxyaromatic substrates by *C. parapsilosis* cells.**

*C. parapsilosis* CLIB214 cells grown in the synthetic media containing a hydroxyaromatic substrate as a sole carbon source at 28 °C till OD_600_ ∼ 1. Substrate consumption was inferred from the absorption spectra (200-350 nm) measured in the media of three parallel cultures (shown in red, blue, and green) after cultivation (t = 17.5, 25.5, and 16 hours for 3-hydroxybenzoate, 4-hydroxybenzoate, and hydroquinone, respectively) as well as in the control medium. Each measurement was performed in three technical replicates. The samples were diluted 2-, 20-, and 5-fold prior analysis of 3-hydroxybenzoate, 4-hydroxybenzoate, and hydroquinone consumption, respectively.

